# A Scalable Staining Strategy for Whole-Brain Connectomics

**DOI:** 10.1101/2023.09.26.558265

**Authors:** Xiaotang Lu, Yuelong Wu, Richard L. Schalek, Yaron Meirovitch, Daniel R. Berger, Jeff W. Lichtman

## Abstract

Mapping the complete synaptic connectivity of a mammalian brain would be transformative, revealing the pathways underlying perception, behavior, and memory. Serial section electron microscopy, via membrane staining using osmium tetroxide, is ideal for visualizing cells and synaptic connections but, in whole brain samples, faces significant challenges related to chemical treatment and volume changes. These issues can adversely affect both the ultrastructural quality and macroscopic tissue integrity. By leveraging time-lapse X-ray imaging and brain proxies, we have developed a 12-step protocol, ODeCO, that effectively infiltrates osmium throughout an entire mouse brain while preserving ultrastructure without any cracks or fragmentation, a necessary prerequisite for constructing the first comprehensive mouse brain connectome.

## Introduction

The wiring diagrams connecting nerve cells underpin brain function and lay bare the extraordinary complexity of the nervous system. Connectomics, a nascent field that studies this connectivity at the level of synapses, primarily employs high-resolution serial section electron microscopy (EM) to obtain synapse-level connectivity maps. This mapping relies on heavy-metal staining of brain tissue. The affinity of heavy-metal stains, most especially osmium tetroxide, for lipids, proteins, and nucleic acids, presents a complete picture of neuronal structure down to the nanometer scale showing the finest neuronal processes and their synapses. Because axons often project long distances, it is difficult to derive functional insights from analysis of small regions of a brain, given that so many nerve cells providing connections to each region are from distant sites and so many neurons in any analyzed region project their output to distant sites. For this reason, whole-brain connectivity maps are required for assigning a function to each neuron and each brain region.

Thanks to rapid advances in automated EM mapping over the past decade, comprehensive connectomes have been reconstructed at the system level in small animals such as worms (*1*) and fruit flies (*2, 3*). However, connectomic mapping has only been feasible for cubic millimeter-scale, or smaller, portions of mammalian brains (*4, 5, 6, 7*). One of the principal obstacles towards larger volumes is the lack of effective osmium staining. Excellent osmium staining has recently been shown in volumes of greater than 10 mm^3^ (*8*) but, even in a small mammal like a mouse, staining whole brains requires the stains to work in samples that are 500 mm^3^. Traditional heavy-metal staining techniques, initially designed for small non-neural tissues, are suboptimal for staining whole mammalian brains. Brain tissue is much more membranous and heterogeneous than most other tissues, making uniform high-contrast staining difficult to achieve. Until a decade ago, no approaches worked in brain volumes as large as one cubic millimeter. The advent of connectomics catalyzed many modifications to traditional techniques. Notably, Hua et al. developed an approach that brought the upper limit of effective en-bloc staining to 1mm^3^ brain punches (*9*). Mikula and Denk went further and demonstrated that a whole mouse brain could be stained with osmium, but their PATCO protocol provided sufficient staining only for myelinated axons (*10*). A few years later, the BROPA protocol, while being able to reliably stain neuropil and myelin, generated suboptimal membrane contrast and large cracks in the stained brain (*11*). The recently developed whole-brain staining protocol by Song et al. (*12*) is a big improvement over previous whole-brain staining, but still creates microbreakages that may extend over a length of 100 μm, and more problematic macroscopic tissue fragmentation in the anterior and posterior poles of the mouse brain. Without a completely intact and well stained brain, a large number of neuronal pathways will be disrupted. We sought to systematically analyze the problems associated with current whole-brain staining approaches to develop an improved strategy. We found that this effort required a better understanding of reaction mechanisms in order to select appropriate reagents for each staining step and a fuller account of the reasons for tissue swelling and shrinkage which causes tissue damage. Moreover, because the final outcome for volumes that are many hundreds of cubic millimeters is only accessible weeks or months after beginning a staining protocol, it was necessary to determine the optimal conditions by leveraging time-lapse X-ray imaging (*13*) and brain proxies. In this way, it was possible to monitor and modify steps in the sample preparation process efficiently without needing to wait for the final result. This approach led to a reliable staining protocol, ODeCO (Osmication-Destaining-Conditioning-Osmication), for adult and juvenile rodent brains without tissue fragmentation, cracks, or weakly stained central areas. The method we describe can be used to stain mouse brains, or potentially even larger mammalian brains, for comprehensive connectivity mapping. ODeCO alleviates perhaps the most serious bottleneck impeding a whole brain connectome in the mouse (*14*), a central thrust of neuroscience research in the coming decade.

## Results

### X-ray microCT reveals the rate of osmium staining and the benefit of maintaining ECS

The electron-based contrast of neural specimens depends on the amount of stain deposited at the sites of interest (mostly membranes) versus the more nonspecific background staining of cytoplasm. As cellular structures have limited heavy metal binding sites, stains that are particularly electron-dense (i.e., have a larger atomic number) are preferable for EM. Osmium tetroxide (OsO_4_), with its large atomic number (Z=76) and ability to polymerize to generate a high density of atoms in a small area, remains the most popular biological stain for EM (*15*), despite being one of the earliest heavy-metal stains (*16*). The nucleus of the osmium atom strongly attracts electrons, providing excellent signals in both scanning and transmission EM imaging. OsO_4_ can bind to various functional groups of biomolecules (e.g., thiols and amines), but more importantly, it imparts high contrast to membrane structures by reacting with unsaturated phospholipids that are the major constituent of the plasmalemma and intracellular membranous structures of cells. This membrane staining is especially important for EM-based connectomic studies because one essential image processing step toward the final reconstruction is partitioning EM images into distinct objects that belong to different cells (i.e., segmentation). Reliably identification of membranes is fundamental for tracing neural processes and pinpointing their termini at synapses, whose internal membrane-bounded synaptic vesicles are also vital for synapse identification.

Although the reaction between OsO_4_ and unsaturated phospholipids is rapid, the diffusion of OsO_4_ varies based on tissue characteristics (*17*). As a result, it is difficult to predict the optimal staining duration for any specific tissue sample. Identifying the best method for staining large tissue volumes could necessitate running hundreds of experiments to determine suitable conditions, with each sample only providing a result after it is fully stained and sectioned, weeks or even months later. We overcame these difficulties by setting up a reaction capsule inside an X-ray computerized tomography (CT) microscope to visualize staining in real time. As shown in Figure 1a, a mouse brain was wrapped in a mesh Nylon bag and sealed in a glass scintillation vial filled with the staining solution. A series of tomographic rotary scans were taken to monitor the progress of the staining (see Methods for the experimental details). The heavy-metal stained parts absorb more X-ray photons than the unstained or under-stained parts, allowing us to visualize the staining front in a large brain sample during the staining process.

**Fig. 1.**
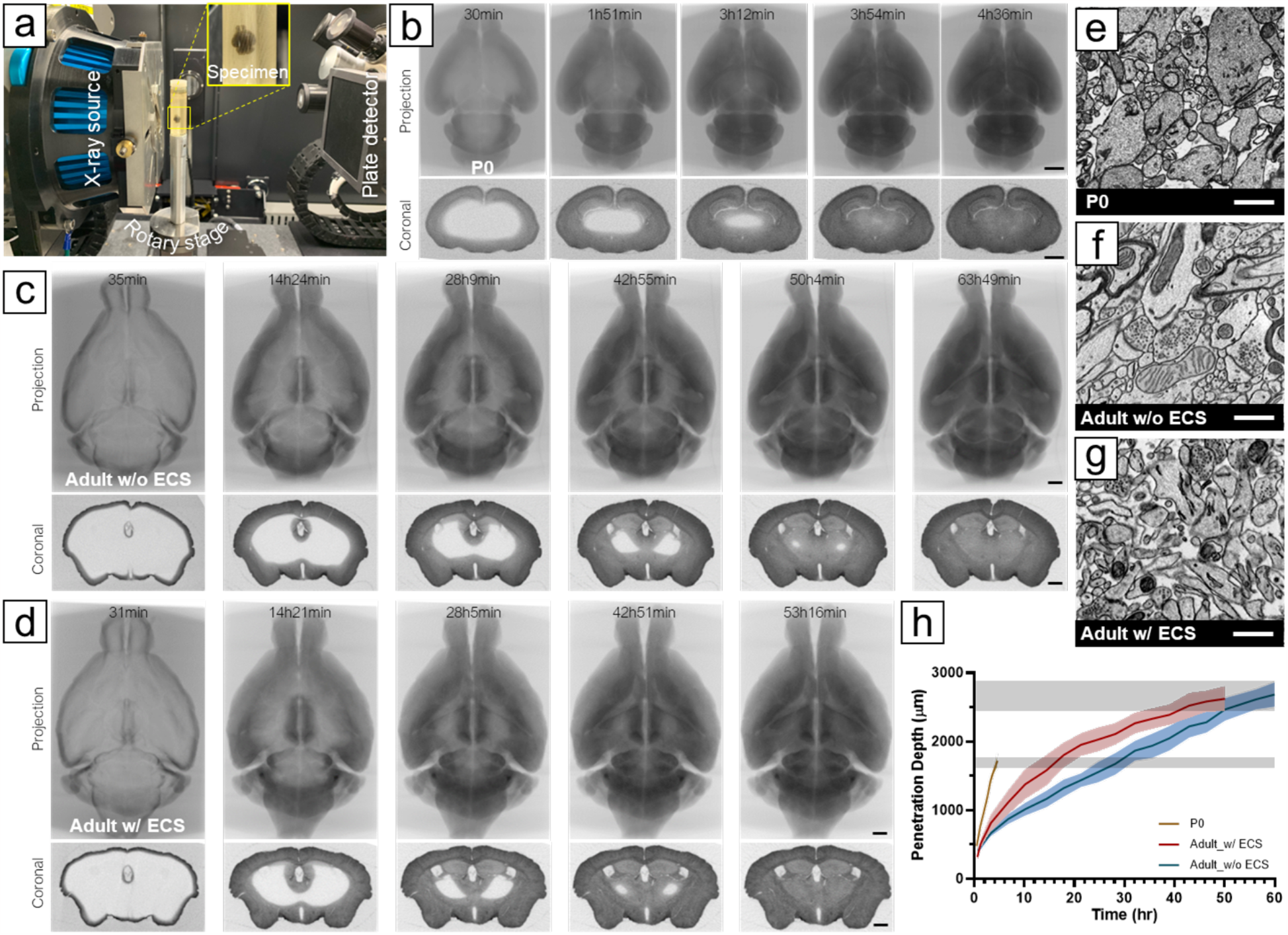
Time-lapse X-ray microCT analysis of osmium staining. (**A**) Experimental setup for time-lapse X-ray imaging, where a mouse brain sealed in a scintillation vial with staining solutions was immobilized on a rotary stage. X-ray photons from the source passed through the sample and were detected by the plate detector. During a tomographic scan, the sample rotated 360°, capturing 401 projections. A series of X-ray microCT projections (upper row) and virtual coronal sections (lower row) obtained during the first osmication were reconstructed from (**B**) a P0 mouse brain (Movie 1), (**C**) an adult mouse brain without preserved ECS (Movie 3), and (**D**) an adult mouse brain with preserved ECS (Movie 4). (**E**) ECS was preserved in P0 mouse brains using conventional perfusion fixation, (**F**) but led to ECS diminishment in adult mouse brains. (**G**) Our modified four-step perfusion fixation method maintained ECS in adult mouse brains. Images were taken from the cortex. (**H**) Plot showing the migration of osmication fronts in a P0 mouse brain, an adult mouse brain without ECS, and an adult mouse brain with preserved ECS during the first osmication. Osmium penetration depths were measured as the shortest distance between the surface and osmication front from different positions in a virtual coronal section near the brain volume’s geometric center at selected time points. The gray bands showed the ending points for staining adult (upper) and neonatal (lower) brains, where unstained regions were no longer observed. Scale bars in b, c, d: 1 mm. Scale bars in e, f, g: 1 μm.

Figure 1b shows a series of average-intensity projection and cross-sectional tomographic images of the first osmication of a neonatal (P0) mouse brain (darker areas have greater osmium impregnation). In the three P0 brains that we imaged with time-lapse, it took 3.4 hours to 4.0 hours for osmium to reach the center of the volume. For adult mouse brains (Figure 1c), completion of the first osmication required about 66 hours. We measured the propagation of the osmication fronts in P0 and adult brains (Figure 1i) and found that the osmication fronts migrated much faster in P0 brains than in adult brains, as indicated by a larger slope. This suggests that the increase in brain size accounts only partially for the greater duration of staining time in adults. While traditional transcardial perfusion with aldehydes preserves abundant extracellular space (ECS) in neonatal mouse brains (∼40%, (*18*)), it appears to cause the ECS to diminish from ∼20% to less than 5% in adult mouse brains (Figure 1e) (*19*). When ECS is diminished, cell membranes are pressed together. We believe osmium bound to cell membranes in the stained parts forms a diffusion barrier for unbound osmium. Without ECS, the only path for unbound osmium to penetrate into the tissue is to pass through this barrier, diffuse through intracellular spaces, and then get through another cell-cell barrier, making staining inefficient for large brains. However, the presence of ECS in P0 brains provides long-range paths for molecules to diffuse, facilitating the staining process.

To improve diffusion efficiency of heavy-metal stains in adult mouse brains, we developed a four-step transcardial perfusion method to preserve the ECS (paper under review). We were able to preserve ECS throughout an adult mouse brain with this perfusion fixation protocol (Figure 1f). As a result, it took about 54 hours to complete the first osmication in an ECS-preserved adult mouse brain (Figure 1d), which was 18% faster than in a mouse brain fixed with the traditional method. While ECS accelerated staining of adult mouse brains, the staining was still slower than in the P0 brains. It is possible that the difference is caused by the fact that in P0 brains there is a greater amount of ECS and reduced tortuosity of the ECS due to sparser neurites and fewer glial processes (*20*), as well as fewer myelinated regions that act as thick barriers to impede osmium diffusion.

### Destaining cytoplasm improves membrane contrast

In large-volume serial section electron microscopy, two rounds of osmium staining are necessary for enhancing the signal and speeding up image acquisition. As osmium reacts with both unsaturated phospholipids and proteins, reducing the amount of osmium bound to cytoplasmic protein will increase the ratio of membrane-bound osmium, leading to improved membrane contrast. More importantly, in whole-brain processing that requires long staining times for both rounds, minimizing osmium binding to cytoplasmic proteins in the first round can improve the efficiency of the subsequent staining, reduce staining duration, and prevent tissue overstaining. We found that without any measures taken to decrease cytoplasmic staining after the first osmication (i.e., using the osmication-conditioning-osmication approach), the second osmication took over three weeks to complete. This not only resulted in all cellular structures appearing dark and indistinguishable (Figure S1), but also made the block too metallic to cut without breakage.

Potassium ferrocyanide (K_4_[Fe(CN)_6_]) is often added together with OsO_4_ to bias osmium to stain membranes rather than cytoplasm (*21*). When mixed, ferrocyanide reduces OsO_4_ to potassium osmate (K_2_[OsO_2_(OH)_4_], which is water soluble but not very stable. It will dissmutate into OsO_4_ and “osmium black” precipitates (i.e., mixture of Os_2_O_5_ and OsO_2_) (*22*). This reaction typically takes several weeks at room temperature to complete; however, in tissue, owing to the abundant presence of unsaturated phospholipids that deplete OsO_4_, the reaction equilibrium shifts to rapidly generate precipitates. Excessive amounts of osmium black produced within a short time will accumulate in the ECS and cytoplasm. In thick tissues, the precipitate builds up close to the surface of the tissue block (∼150 μm deep, Figure S2), acting as a barrier for osmium stains to reach the interior of the specimen. Therefore, this approach (i.e., OsO_4_ and ferrocyanide are applied simultaneously) can only be applied to stain tissue thinner than ∼300 μm.

To solve the problem of the accumulation of osmium black as a barrier to osmium penetration into deeper structures, several strategies have been attempted but are insufficient for routine whole-brain staining. In the BROPA protocol (*11*), formamide is added with the ferrocyanide-reduced osmium to prevent osmium black precipitation by increasing its solubility. But because formamide removes some aldehyde-protein crosslinks (*23*), it leads to adverse side effects, including protein loss and tissue distortion due to tissue softening (Figure S3), especially when the staining protocol is long. The ORTO protocol (*9*) sought to solve the osmium black precipitation problem by separating the first osmication and the ferrocyanide steps. The idea behind this approach is that OsO_4_ is not the actual membrane stain, but osmium black is. By allowing OsO_4_ to penetrate the entire sample and then applying the ferrocyanide, they could induce the reaction that produces osmium black to occur locally without causing overaccumulation issues. They also argued that, because the osmate ester formed in membrane bilayers from the reaction between OsO_4_ and unsaturated phospholipids could be easily hydrolyzed and washed off, it is essential to avoid rinsing the sample between the first osmication step and the ferrocyanide step to preserve as much OsO_4_ in the tissue as a substrate for producing osmium black. While the ORTO method is successful in staining 1 mm-thick brain tissues, it has not been proven effective for larger tissues.

We have found that thorough rinsing between the first osmication and the ferrocyanide steps increases osmium penetration into large volumes. Contrary to our expectation based on the ORTO work, rinsing did not promote a significant release of osmium through hydrolysis, as the extent of X-ray attenuation was only slightly reduced (Figure S4). We believe this is because the osmylation of unsaturated phospholipids takes place in the hydrophobic interior of membrane bilayers, making the hydrolysis of osmate esters rare without a catalyst (*24, 25*). Rinsing likely only washed out the unbound and loosely bound osmium in the cytosol, leaving the more prevalent covalently bound osmium abundant in membranes unaffected. The ORTO work also implied that the membrane contrast would significantly increase after applying ferrocyanide due to the accumulation of osmium black in membrane bilayers. We thus expected that the X-ray attenuation would increase accordingly. In contrast, we found the X-ray signal progressively diminished during the ferrocyanide step in the P0 mouse brain (Figure 2a), indicating that the amount of osmium in the tissue was being lowered by a reaction with ferrocyanide. This meant that potassium ferrocyanide was destaining the tissue. To understand how separation of the osmication and the ferrocyanide steps could improve membrane contrast and which portion of the tissue osmium was being removed by ferrocyanide, we created three simple test blocks for OsO_4_ to stain: an agarose-only gel, an agarose gel infused with unsaturated phospholipids, and an agarose gel infused with serum proteins (see experimental details in Figure S5). After osmication, the phospholipid- and the protein-containing gels turned dark, whereas the agarose-only gel appeared unstained. The staining of all three samples was not altered by rinsing in buffer. These results show that OsO_4_ can form covalent bonds with unsaturated phospholipids and proteins, but not with carbohydrates like agarose. Post-rinsing, we added potassium ferrocyanide to the three gels. The result was striking: the phospholipid gel remained black, while the protein gel was decolorized. It suggests that ferrocyanide does not affect osmium bound to membrane phospholipids, but can dissociate osmium bound to proteins. Ferrocyanide therefore leads to reduced X-ray attenuation but nonetheless enhances membrane contrast by lowering the background staining from cytoplasmic proteins. Based on these findings, we believe that membrane staining is not due to osmium black (i.e., previously suggested as the product of ferrocyanide treatment, (*9*)) but rather to OsO_4_. By separating the addition of OsO_4_ from that of ferrocyanide, we in essence have changed the role of ferrocyanide from a stain additive to a stain remover. Based on this finding, we suggest adding a thorough rinse before introducing ferrocyanide to avoid the reaction with excessive free-floating OsO_4_ (i.e., ferrocyanide acts as a stain additive) that causes the build-up of osmium black. Inserting a washing forces ferrocyanide to react with protein-bound osmium (i.e., ferrocyanide acts as a stain remover), which is crucial for scaling up the staining from 1mm^3^ brain tissues to larger brain volumes. However, from the time-lapse X-ray imaging of the destaining of P0 mouse brains (∼90 mm^3^), we noticed that destaining using ferrocyanide progressed rather slowly (Figure 2a), taking at least five times longer than the OsO_4_ staining. This observation suggested that destaining an adult mouse brain (∼500 mm^3^) could take more than 15 days, raising the question of whether ferrocyanide is the best choice for destaining an adult mouse brain.

**Fig. 2.**
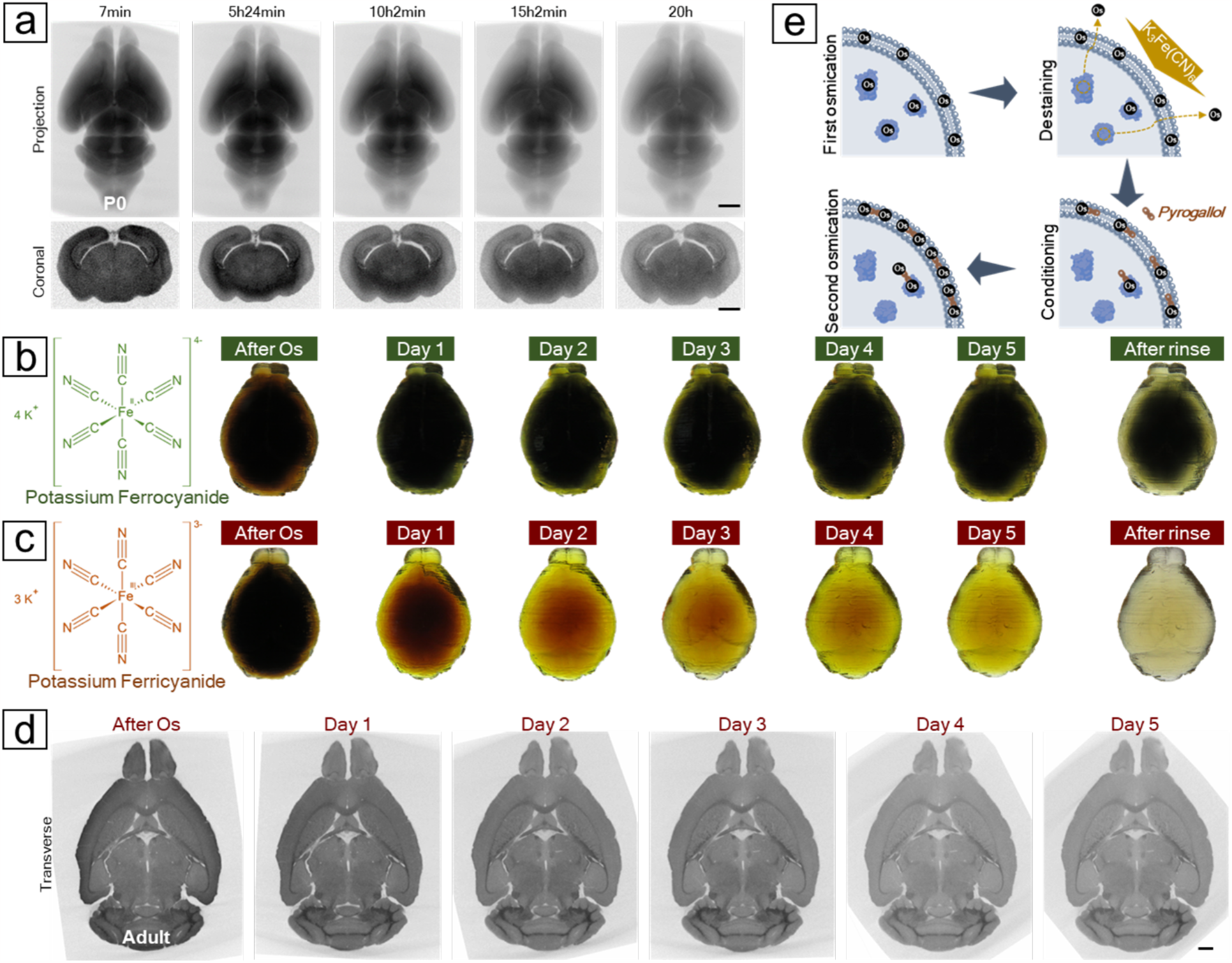
Destaining the osmicated brain to remove excessive cytosolic osmium. (**A**) Time-lapse X-ray microCT revealed a progressive decrease in X-ray attenuation from an osmicated P0 mouse brain after the application of potassium ferrocyanide. (**B**) Destaining a protein-infused mouse brain proxy with ferrocyanide showed that after five days treatment, only the superficial layer was efficiently destained. (**C**) The brain proxy treated with ferricyanide became clear after five days of treatment. As ferrocyanide turned dark green and ferricyanide remained yellow during the reaction, a thorough rinse was applied after five days to reveal the actual color of the brain proxies. **(D)** A virtual transverse section from the X-ray microCT reconstruction of a mouse brain treated with ferricyanide displayed a decrease in X-ray attenuation, especially in gray matter areas, which enhanced the contrast of the white matter tracts over the course of the reaction. (**E**) A reaction mechanism was proposed based on the X-ray observations and the brain proxy tests. Initial osmication with OsO_4_ stains both cell membranes and cytoplasm. Cytoplasmic osmium bound to proteins can be stripped and washed off using ferricyanide, while the remaining osmium anchors multidentate ligands, such as pyrogallol, which provide more binding sites for the second round of osmium binding. Scale bars: 1 mm.

We found that potassium ferricyanide (K_3_Fe(CN)_6_) is more efficient than potassium ferrocyanide in removing cytosolic osmium. Since the majority of X-ray attenuation is caused by membrane-bound osmium rather than protein-bound osmium, tracking the reaction kinetics of the destaining step and determining the endpoint with X-ray microCT becomes difficult. To compare different destaining reagents, we again used protein-infused agarose gels shaped like mouse brains (see Methods for creating dummy brains using 3D printing) to resemble lipid-free brains containing only proteins. By excluding interference from membrane-bound osmium, we were able to directly observe the destaining progress by monitoring the decolorization of the osmicated gels. We found that potassium ferricyanide is a more efficient stain remover (Figure 2b-d). The osmicated dummy brain returned to transparency in just five days after treatment with ferricyanide; in comparison, only the superficial layer of the dummy brain became clear after treatment with ferrocyanide for the same duration. We believe that both ferrocyanide and ferricyanide can directly coordinate with protein-bound osmium through the nitrogen atoms of cyano groups, making these osmium more soluble and eventually releasing them from protein binding. This process is less likely to occur with phospholipid-bound osmium because the osmium residing in the hydrophobic interior of the membrane bilayer is more densely packed, making it less accessible to large anions such as ferrocyanide and ferricyanide. An early study showed that free cyanides (CN^-^), which are not complexed with Fe(II) or Fe(III) and thus much smaller, are a stronger stain remover and can even remove lipid-bound osmium (*26*). Comparing ferrocyanide with ferricyanide, we believe two main factors contribute to the slow kinetics of ferrocyanide. Ferrocyanide ([Fe(CN)_6_]^4−^) has a higher charge than ferricyanides ([Fe(CN)_6_]^3−^) which results in stronger electrostatic repulsion when coordinating with protein-bound osmium. Second, water molecules are more closely bound to ferrocyanide due to its higher charge, making ferrocyanide less diffusive than ferricyanide. (*27*)Additionally, iron in ferricyanide is in its highest oxidation state and will not induce a redox reaction with unbound osmium, circumventing the overaccumulation of osmium black even in the very large brain volume where the interior free-floating osmium might not be thoroughly removed.

Ferrocyanide also induced a considerable 30.9%±2.3% (n=5) volume expansion, significantly larger than the 12.3%±1.9% (n=5) observed after ferricyanide treatment. This greater volume expansion increased the likelihood of membrane breaks forming during this step and cracks forming during subsequent procedures due to heightened brain shrinkage. As a result, in our ODeCO protocol, we substituted ferrocyanide with ferricyanide for more effective, rapid, and minimally size-altering destaining.

### Minimizing volume changes during the pyrogallol conditioning step

The first osmication likely saturates most binding sites because tissue X-ray attenuation reaches a plateau. Since imaging speed is particularly important for large volume serial section electron microscopy, adding more osmium is necessary to speed up image acquisition (e.g., by providing more signal). One way to increase the osmium signal further is to use bound osmium to initiate the formation of additional osmium complexes. This idea of using osmium as the substrate for further osmium deposition is the basis of several popular staining protocols ((*28*) (*29*) (*9*)). In these enhanced osmium staining protocols, a multidentate osmophilic ligand is added after the first osmication to serve as an anchor for additional osmium. Thiocarbohydrazide (TCH) is a widely used ligand for such purposes. In OTO, rOTO and ORTO, TCH gives rise to high membrane contrast. However, when applied to large tissue volumes, TCH produces two artifacts: cracks (Figure 3a) and membrane breaks (Figure 3b). The reaction of TCH is accompanied by the liberation of gaseous nitrogen oxides (NO_x_) and sulfur oxides (SO_x_). Due to the interplay between diffusion and aggregation, these gas molecules may cluster into bubbles before finding their way to the tissue surface. Thicker tissues (i.e., more gas molecules and longer diffusion path) tend to incur more cracks. Another phenomenon that might contribute to the formation of cracks is substantial volume expansion during the TCH step. Due to the low solubility of TCH at room temperature, it is typically applied at elevated temperatures (i.e., 40°C (*9*) to 50°C (*10*)) to prevent its precipitation. However, the high temperature leads to a nearly 80% expansion of tissue volume in our experiments. Such a large volume expansion is difficult for the osmicated tissues to accommodate which gives rise to cracks and membrane breaks (perhaps due to a decrease in the elasticity of metalized membranes). These macroscopic cracks and more nanoscale membrane gaps pose problems for accurate automated cell segmentation.

**Fig. 3.**
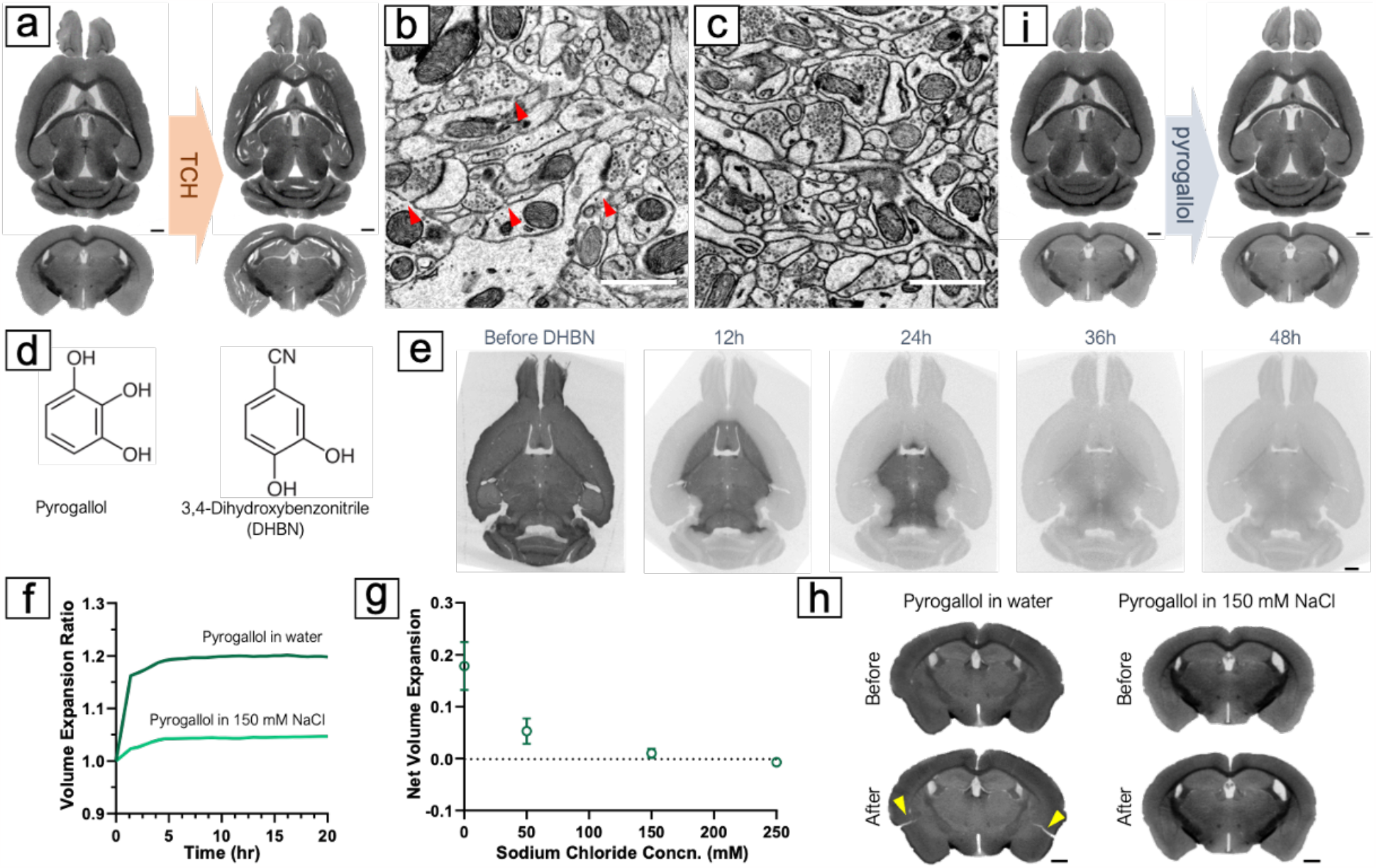
Conditioning the brain with pyrogallol for the second osmium binding. (**A**) Numerous cracks appeared in the brain after reacting with TCH. (**B**) Membrane breaks (indicated by red arrows) were observed in the 1mm brain slabs treated with TCH, likely due to significant but gradual volume expansion during the TCH reaction at 40°C. (**C**) No observable membrane breaks were found in the 1mm brain tissues treated with pyrogallol, and membranes appeared thicker and with higher contrast after reaction with pyrogallol. (**D**) Comparison of chemical structures of pyrogallol vs. DHBN. (**E**) Treatment with DHBN resulted in a decrease in X-ray attenuation in the osmicated mouse brain. X-ray microCT revealed it took approximately 48 hours for DHBN to completely destain an adult mouse brain. (**F**) Applying pyrogallol in water led to an immediate 20% volume expansion of the brain, while using 150 mM NaCl reduced the volume change. (**G**) Comparing different concentrations of NaCl solution as a reaction medium showed that using 150 mM NaCl was optimal, which matched the osmolarity of the cacodylate buffer used in previous procedures. (**H**) The volume expansion during the pyrogallol conditioning step without adjusting the reaction medium caused cracking of the brain (indicated by yellow arrows), while using 150 mM NaCl solution with pyrogallol resulted in no noticeable cracks. (**I**) No crack was observed in the mouse brains after applying pyrogallol with adjusted media osmolarity. Scale bars in a, e, h, i: 1 mm. Scale bars in b, c: 1 μm.

Pyrogallol was proposed to replace TCH for whole mouse brain staining in the BROPA protocol by Mikula and Denk (*11*) due to its increased solubility. Using pyrogallol eliminates the need for high temperatures, it also produces no gas resulting in much less tissue damage. In our test with small samples of 1mm thick, no cracks or membrane breaks were observed, and membranes developed excellent contrast (Figure 3c). However, when moving to whole-brain staining, we faced a technical challenge: as an organic compound, pyrogallol has much lower X-ray absorbance than metals, making it difficult to monitor the process of this step using X-ray microCT. To determine the optimal reaction time, we used 3,4-dihydroxybenzonitrile (DHBN), a chemical analog of pyrogallol, to treat osmicated mouse brains. Like pyrogallol, DHBN binds to osmium through its hydroxyl groups. But with one of the three hydroxyl groups replaced by a cyano group (Figure 3d), DHBN can release membrane-bound osmium into the aqueous solution (see discussion about cyanides above). This significantly decreases tissue X-ray attenuation (Figure 3e), making it unsuitable for staining but useful for inferring reaction duration for pyrogallol. We found that it took approximately 48 hours for DHBN to destain an osmicated mouse brain, and inferred that a similar amount of time would be appropriate for pyrogallol to penetrate the entire brain to bind to tissue-bound osmium.

Despite the many advantages of pyrogallol over TCH for conditioning large brain tissues, one drawback of using pyrogallol is that it is acidic and cannot be used with the mildly basic sodium cacodylate buffer at pH 7.4. In an acidic or neutral milieu, pyrogallol forms stable complexes with osmium atoms (*30*) setting itself up as the substrate for the next round of osmium binding. The BROPA protocol used an unbuffered aqueous solution of pyrogallol. However, we found that a sudden change in the reaction medium from cacodylate buffer to water resulted in an immediate 17.8%±4.6% (n=3) volume expansion of the brain (Figure 3f), giving rise to cracks (Figure 3h). This is because pyrogallol undergoes almost no dissociation to pyrogallic acid in water (pKa = 9.01), so the ion concentration of the pyrogallol aqueous solution is much lower than that of the buffer with the same molarity. This drop in ion concentration leads to the sudden tissue expansion and causes tissue cracking. To solve this problem, we added sodium chloride (NaCl) to adjust the ion concentration of the pyrogallol solution to match that of the cacodylate buffer used in the previous rinsing. Our results showed that this adjustment resulted in minimal volume change during pyrogallol impregnation of mouse brains and no cracks (Figure 3i).

### Using Sodium Chloride instead of Sodium Cacodylate for the second osmication

Upon changing the reaction medium to sodium chloride during the pyrogallol conditioning step, we were presented with two options: continue using sodium chloride or revert to sodium cacodylate. We tested both and discovered that sodium chloride considerably shortened the staining time for the second osmication. Using sodium cacodylate, the second osmication took over 14 days to complete (Figure S6), while sodium chloride reduced this duration to just 6 days. It has been hypothesized that cacodylate anions may coordinate with osmium (*12*), potentially increasing the hydrodynamic radius of osmium tetroxide and thereby reducing its diffusivity. Coordination with large anions may also introduce steric hindrance, which could decrease the reactivity of aqueous osmium with the densely packed osmium in the tissue. Nevertheless, we noticed that substituting sodium cacodylate with sodium chloride in the first osmication adversely affected staining quality, as the membranous structures were inadequately stained (Figure S7). Thus, we still recommend using sodium cacodylate for the first osmication. Since the second osmication process takes considerably longer than the first and does not involve a subsequent destaining step, we lowered the reaction temperature for this step to prevent overstaining of the cytosol.

### Corrective strategies prevent artifacts over the course of sample processing

Staining protocols are commonly associated with volume changes. While these changes are usually negligible in thin tissues and not well documented in previous studies, they become more evident and can give rise to artifacts when scaling up to whole mouse brains. We observed that small membrane breaks often resulted from gradual yet notable volume expansion, whereas larger tissue cracks were caused by abrupt volume changes, either shrinkage or expansion. To address these issues, we monitored volume changes in mouse brains using X-ray microCT reconstructions acquired during or after each step (see Methods for technical details). By tracking volume changes, pinpointing when cracks occurred, and investigating potential causes, we attempted to develop a set of strategies to minimize the impact of problematic volume changes.

In previous staining studies, the reaction medium is often changed from buffer to water after the first osmication. For example, the ORTO protocol switches to water after the ferrocyanide step (*9*). However, we found that the transition from buffer to water induced severe damage to the large-volume specimens. In particular, when we transferred an osmicated P0 brain from 150 mM cacodylate buffer to water following the destaining step, the brain disintegrated as its volume rapidly swelled within just a few minutes (Figure S8a). We believe the sudden drop in ion concentration in the rinsing solution causes the cross-linked proteinaceous network of the tissue to swell abruptly as water molecules insert themselves into the existing structure (*31*). A simple test supported this hypothesis: when we moved a chemically fixed, unstained neonatal mouse brain and an adult mouse cerebellum from cacodylate buffer to water, an immediate and perceptible volume expansion (approximately 15%) occurred (Figure S8b). Although this expansion can be accommodated by the elasticity of unstained tissue, samples stained with heavy metals become brittle and lose their elastic properties, resulting in disintegration. Previous whole-brain staining protocols have used water transitions as rinses before (*12*) and after (*11*) the pyrogallol step. In these protocols membrane breaks, fissures, fragility, and disintegration have been reported previously and confirmed by us (Figure S9). To prevent this issue, we maintained the osmolarity of the reaction medium through the staining and rinsing steps up to the final graded dehydration steps. This strategy prevented the dramatic tissue volume changes associated with solution-to-water transitions and avoided cracks and membrane breaks.

After the second osmication, the brain samples were rinsed with sodium chloride before undergoing dehydration for resin infiltration and embedding. Tissue dehydration typically causes shrinkage (*31*). Traditional dehydration methods involve a stepwise exchange of solvents at concentrations of 25%, 50%, 75%, and 100%. Due to the large concentration differences between the steps, sudden shrinkage occurs at each solvent change. To avoid artifacts resulting from abrupt volume shrinkage, we adopted a gradual dehydration strategy instead of the stepwise approach. Our device was modified on a previous design (*32*) that allowed for rate-controllable solution exchange (see Figure S10 for technical details). We used a syringe pump to slowly inject acetonitrile into a container holding the specimen in sodium chloride at a preset flow rate. The container was approximately 100 times the volume of the mouse brain. As acetonitrile was added, the solution continuously mixed with a spinning stirrer bar, and excess solution automatically drained from an overflow hole. Once the acetonitrile concentration reached 25%, we replaced the container’s solution with a mixture of 25% acetonitrile and 75% water to prevent sodium chloride precipitation. We then resumed acetonitrile injection until its concentration reached 99%. Subsequently, the specimen was transferred to pure acetonitrile for final rinsing before being infiltrated with a diluted resin solution. We found that a threshold slow dehydration rate, dependent on the sample volume, was crucial to prevent crack formation. For example, a 6-hour gradual dehydration resulted in cracks in a P0 mouse brain (Figure 4a), whereas 20-hour dehydration prevented them (Figure 4b). For an adult mouse brain, 96-hour dehydration was necessary to avoid cracks (Figure 4c-d). Despite introducing gradual dehydration, we could not entirely prevent volume shrinkage. There was still 11.9±0.5% (n=4) of volume shrinkage. Instead of cracking, this shrinkage of brain parenchyma only led to expansion of the tissue-free ventricles (Figure 4d). We think this is a tolerable artifact for whole-brain imaging.

**Fig. 4.**
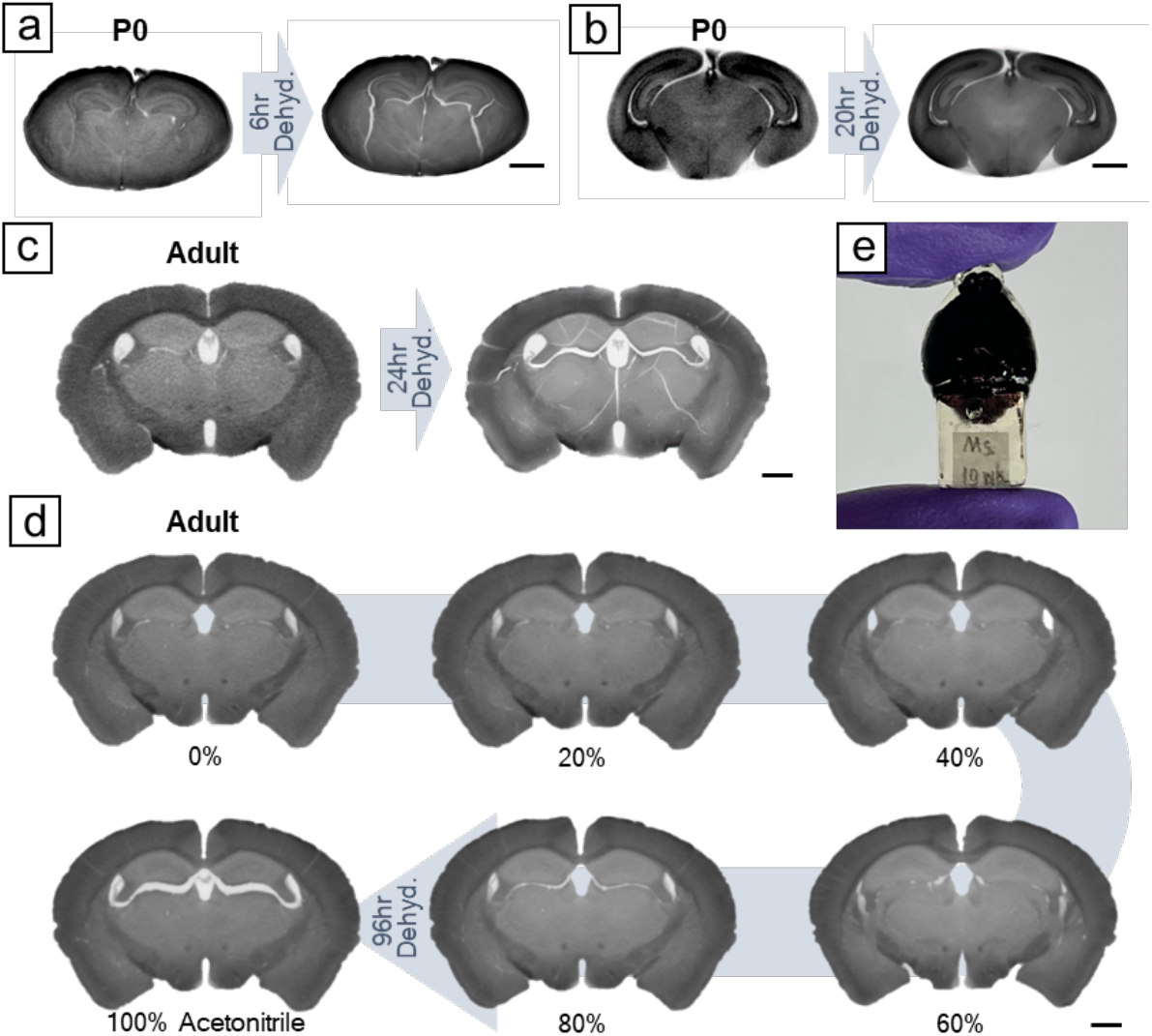
Gradual dehydration prevents cracking during tissue dehydration. (A) For P0 mouse brains, a 6-hour gradual dehydration caused cracking, (**B**) while no noticeable cracks were observed in a 20-hour dehydration. (**C**) For adult mouse brains, a 24-hour gradual dehydration resulted in tissue cracking. (**D**) Extending dehydration to 96 hours prevented tissue cracking. There was still volume shrinkage in a 96-hour dehydration, which became noticeable after acetonitrile achieved over 80%. Instead of cracking, this shrinkage led to the enlargement of brain ventricles. **(E)** Photo of a resin-embedded adult mouse brain with a gripping handle on the bottom. Scale bars: 1 mm.

Previous whole-brain staining attempts employed Spurr’s resin for tissue embedding and used block-face scanning electron microscopy for imaging (*10–12*). The lower viscosity of Spurr’s resin, compared to the more commonly used Epon epoxy resin, allowed for easier and more complete infiltration into large brain samples. The cut-away sections are not imaged in the block face scanning electron microscopy technique so the quality of the cut-away sections is not important. Since we plan to cut the whole brain sample into semi-thin serial sections (∼20,000 sub-micron sections) and image them with multibeam scanning electron microscopy and ion beam milling (*33*), we tested the cutting properties of Spurr’s embedded mouse brains and found that the sections featured many small tears and accumulated debris. Therefore, we tested other resins and found that UltraBed resin, a modified formula of Spurr’s, allowed for rapid infiltration at room temperature without premature polymerization, and demonstrated improved cutting properties compared to Spurr’s, with significantly fewer cutting artifacts.

We further modified the resin embedding methods to prevent microcracking, a phenomenon we only observed in whole-brain specimens. We noticed that when using off-the-shelf rectangular molds, microcracking occurred in the tissue after cutting open the resin-embedded brains for tissue screening. We believe that microcracking is caused by the differences in cure shrinkage between the resin-infiltrated metal-stained tissue and pure resin surrounding the tissue, leading to a buildup of stress in the tissue during resin curing. When the tissue is suddenly cut open, the internal residual stress is no longer in equilibrium, causing microcracks to nucleate at the pure resin/tissue interface and propagate within the tissue, a phenomenon similar to that observed in glass-ceramic composites (*34*). To address this issue, we first developed customized silicone molds for whole-brain resin embedding (Figure S11 and see Methods for technical details). This mold reduced the volume of the resin surrounding the tissue, thereby inducing less stress. Along with the embedding mold, we also modified the heating and cooling process for resin curing. Instead of placing and removing samples from an already heated oven as is typically done, we programmed the oven to gradually increase the temperature from room temperature to 70°C over a 20-hour ramp, maintain the 70°C temperature for 24 hours to cure the resin, and then cool down over another 20-hour ramp to room temperature. This helped reduce the potential buildup of internal residual stress. After implementing these modifications, we no longer encountered the microcracking issue (Figure S12).

### Assessment of ultrastructure with the ODeCO protocol

To verify the feasibility of the ODeCO protocol, we applied it to stain 1mm-thick brain slices, whole P0 mouse brains (∼5.6mm×8.9mm×3.5mm), and adult mouse brains (∼10.5mm×16.5mm×6.3mm; refer to Methods for protocol details). Staining uniformity and the absence of tissue artifacts were first confirmed by X-ray microCT at micrometer resolution. To examine the ultrastructural quality, we examined the ODeCO-prepared P0 and adult mouse brains at various brain regions (Figure 5) by tape collection-based serial section electron microscopy (see Methods; (*35*)). The imaged regions were largely completely free of artifacts including cracks, membrane breaks, and heavy-metal precipitates. High-contrast staining was confirmed throughout the entire volume and there was no noticeable attenuation of staining quality with depth (a link to access all the screening data: https://lichtman.rc.fas.harvard.edu/whole_brain_staining/).

**Fig. 5.**
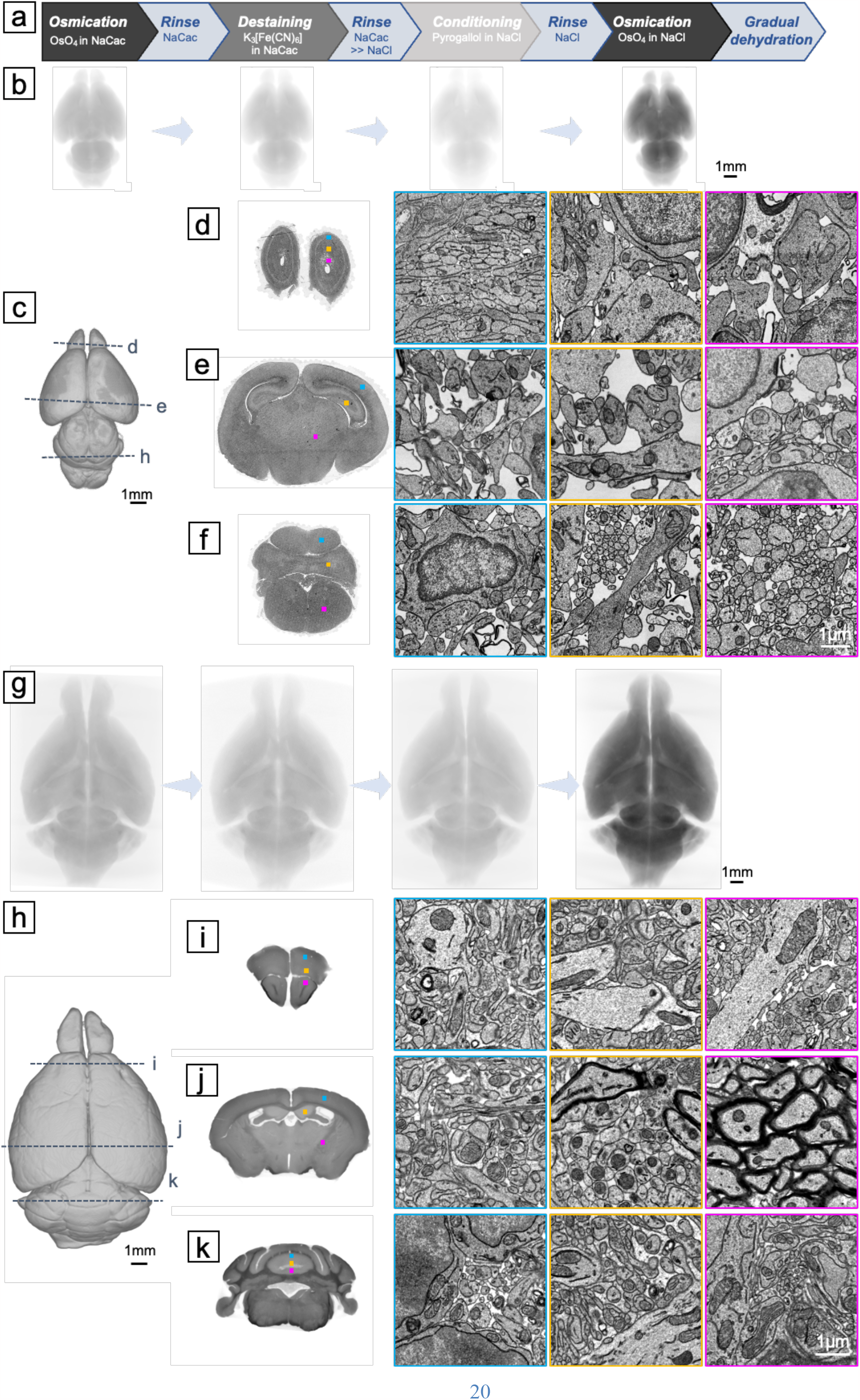
Ultrastructural assessment of the ODeCO protocol. (**A**) Illustration of the ODeCO protocol workflow. (**B**) X-ray microCT projections of a P0 mouse brain throughout the ODeCO procedure. The color was normalized to the last microCT scan. The progression in X-ray attenuation intensity showed the changes of osmium density in the tissue during the ODeCO procedure. (**C**) Reconstruction of an ODeCO-stained and resin-embedded P0 mouse brain, demonstrating the tissue remained intact and free of crack (also see Movie 5). (**D-F**) Ultrathin coronal sections were collected from various brain regions and imaged at 4nm×4nm. More sections at full resolution are accessible through the provided link. Magnified views of selected regions, as indicated by the corresponding colored frames and squares on the coronal sections, showed high membrane contrast staining achieved consistently across the neonatal tissue volume. (**G**) X-ray microCT projections of an adult mouse brain during the ODeCO procedure. (**H**) Reconstruction of an ODeCO-stained and resin-embedded adult mouse brain. (**I-K**) Virtual coronal sections from X-ray microCT affirmed that staining was uniform through the volume, with no evidence of cracking or fragmentation in the anterior, central, and posterior of the brain (also see Movie 6). Similarly, magnified views of certain regions, indicated by the corresponding colored frames and squares on the virtual coronal sections, showed that high membrane contrast staining was attained throughout the adult tissue volume.

We also sectioned 109 consecutive ∼35nm full-size coronal sections (∼5.16mm×3.67mm) from an ODeCO-stained P0 mouse brain. Using a multibeam scanning electron microscope, we acquired at 4nm×4nm an image stack covering layers II/III through VI of M2 (i.e., part of motor cortex; ∼180μm×303μm×4μm) from the serial sections (Figure S13). Qualitatively, we found it was easy for human annotators to segment and trace even the finest neurites in the aligned image stack. To examine if the quality of this dataset was sufficient for automated segmentation, we implemented a lightweight segmentation pipeline to reconstruct cells and their processes (see Supporting Information for technical details). We rarely detected merge errors (n=89 or 0.00043 merges/μm^3^) where two independent objects were erroneously connected. More commonly we observed split errors, where one object was divided into multiple objects. We picked three random sub-volumes of 4μm×4μm×1μm to measure the split error rate. There were on average 0.74 splits/μm^3^. To compare these merge and split error rates to other datasets, we added two sub-volumes that contained one merge error and many split errors in each. We identified 0.22 errors/object in our sub-volumes. This error rate was slightly lower than the error rates (∼0.3 errors/object) found in an adult ECS-preserved mouse brain dataset (*36*). We also measured the variation of information (VI) which compared the pixel differences between automated segmentation and ground truth segmentation from human annotators (*37*). The VI of our auto-segmentation was 0.1144, which was close to the estimate of human accuracy (*38*). All of these results suggest that the quality of the whole brain staining was quite good for automated segmentation.

## Discussion

To meet the needs of increasingly larger connectomic projects, especially the whole mouse brain project, we developed ODeCO, a procedure that is able to stain and embed whole mouse brains for EM imaging. ODeCO is comprised of two rounds of osmication as reflected in its name: O, the first osmication in which OsO_4_ binds primarily to unsaturated phospholipids and proteins to provide the first layer of staining; De, the destaining step that involves application of potassium ferricyanide to selectively remove osmium bound to cytosolic proteins; C, the conditioning step that involves application of pyrogallol to prime the tissue for another round of osmium binding; and finally O, the second osmication to enhance the pre-existing osmium stain.

Using X-ray microCT and brain proxies, we developed a comprehensive understanding of the mechanisms of staining and the conditions required to achieve uniform results without artifacts in ∼500 mm^3^ brain specimens. We discovered that preserving extracellular space in adult mouse brains improved staining efficiency. Our investigation also led to a reinterpretation of the role of potassium ferrocyanide in the process. Ferrocyanide has been considered as a helper reagent that reduces osmium tetroxide and increases its specificity for membrane staining. However, this role for potassium ferrocyanide only applies to the situation where it is added together with osmium tetroxide, as in the rOTO protocol, which however is not suitable for large specimens. We found that, when ferrocyanide is added separately after osmium tetroxide staining, it actually acts as a destaining reagent. We believe that it enhances membrane contrast not by directly staining membranes but by reducing cytosolic osmium staining. To achieve high membrane contrast through the entire brain volume, sufficient destaining is needed to allow the cytosolic osmium located in the interior to be removed. Based on this understanding, we found that potassium ferrocyanide is not an efficient destaining reagent for adult mouse brains, as the process is too slow and insufficient. Potassium ferricyanide, on the other hand, exhibits significantly better reaction kinetics, making it a more suitable choice for large brain tissues.

Beyond staining chemistry, we also rigorously investigated the volume changes in specimens and their connection to tissue cracking during the staining, rinsing, and dehydration steps. The volume changes have not been discussed in depth before, as they do not typically cause noticeable artifacts in small specimens. However, we found that sudden volume expansion during the solution-to-water transition and shrinkage during the stepwise dehydration were the primary causes of macroscopic cracks in brain specimens. To address these issues, we used solutions with consistent osmolarity throughout the staining and rinsing steps and implemented a gradual solution exchange system for dehydrating the osmicated brain samples at an appropriate rate depending on tissue volume. We found that these modifications helped prevent abrupt volume changes and prevented the formation of cracks. To improve the cutting properties of such large brain tissue for serial sectioning, we employed an optimized embedding method for the whole-brain samples utilizing a previously rarely used resin (i.e., UltraBed low-viscosity epoxy), customized brain molds, and ramped heating and cooling to reduce internal stress build-up in the resin block.

There is a long history of attempts to stain whole mammalian brains with osmium. Sixty years ago, Palay and colleagues (*39*) perfused osmium tetroxide into a rat brain via the vasculature. In our attempts to replicate this approach in mouse brains, we found that achieving uniform staining was problematic, as multiple brain regions were poorly infiltrated with osmium. Connectomics however has spurred a number of newer staining approaches (*8, 9, 29, 40, 41*). While most of these were not designed for volumes as large as a mouse brain, the insights gained from these investigations have proven invaluable for scaling methods for larger volumes. In particular, the introduction of two rounds of osmication for enhanced membrane contrast (*29, 40*), the separation of osmium and ferrocyanide treatments to increase the penetration depth of staining (*9*), and the preservation of ECS to improve staining efficiency (*36*) have all advanced the field. Moreover, the recent attempts at staining a whole mouse brain have replaced TCH with pyrogallol for tissue conditioning (*11, 12*) and employed microCT X-ray monitoring of the staining process (*42*), greatly improving large volume staining and its analysis. Nevertheless, multiple challenges persist because, for a sample the size of a mouse brain, improving whole-brain staining quality often has deleterious effects on tissue integrity. The slight volume changes, that heavy-metal staining and resin embedding give rise to, are significantly magnified in volumes of hundreds of cubic millimeters, leading to nanoscale membrane breaks, microscopic fissures, and macroscopic cracks in the metalized tissue. Starting from the foundation of previous staining protocols, we developed ODeCO, a method that minimizes volume changes and thus allows all staining steps to be scaled with time. As evidence, ODeCO has been successfully applied to both small and large brain specimens: staining a 1mm-thick brain slice took ∼70 hours, a neonatal mouse brain ∼8 days, and an adult mouse brain ∼6.4 weeks. Extrapolating from this, we think a marmoset brain could also be stained this way although it would likely take nearly two years. Even if one could stain tissues as large as a primate brain, the methods to section and image such large volumes would likely require novel solutions. In summary, ODeCO overcomes the remaining challenges of existing staining methods, making it a suitable method for mapping the mouse brain connectome.

## Acknowledgments

We would like to thank Dr. JoAnn Buchanan for sharing her experience using ferricyanide in tissue staining. X.L. is supported by the NIH BRAIN Initiative award K99MH128891. This research was also supported by NIH grants U19NS104653 and UG3 MH123386.

## Author contributions

XL was responsible for the method development and experimental execution. YW handled the analysis of X-ray microCT, stitching and alignment of EM images, as well as the creation of 3D printed brain models. RLS provided support in sample microtomy and both EM and X-ray imaging. YM conducted automatic segmentation, while DRB contributed to 3D reconstruction rendering. The manuscript was drafted by XL and JWL, with valuable input from all other authors.

## Competing interests

Authors declare that they have no competing interests.

## Data and materials availability

All data are available in the main text or the supplementary materials. Additional data is accessible through the link https://lichtman.rc.fas.harvard.edu/whole_brain_staining/

## Notes

### Competing Interest Statement

The authors have declared no competing interest.

