## Supplemental Materials for "A Scalable Staining Strategy for Whole-Brain Connectomics"

### Materials and Methods

Materials

**Table S1.** Chemical reagents and algorithms used in this work.

| **Reagents** | **Suppliers** | **Catalog Number** |
|---|---|---|
| Sodium cacodylate trihydrate | Electron Microscopy Sciences | #12310 |
| Calcium chloride solution, 1M | Sigma-Aldrich | #21115 |
| Magnesium sulfate solution, 1M | Sigma-Aldrich | #M3409 |
| 32% paraformaldehyde solution | Electron Microscopy Sciences | #15714 |
| 25% glutaraldehyde solution | Electron Microscopy Sciences | #16220 |
| 4% Osmium tetroxide solution | Electron Microscopy Sciences | #19190 |
| Potassium ferrocyanide (potassium hexacyanoferrate(II) trihydrate | Sigma-Aldrich | #60279 |
| Potassium Ferricyanide (Potassium hexacyanoferrate(III)) | Sigma-Aldrich | #60299 |
| Pyrogallol | Sigma-Aldrich | #16040 |
| Acetonitrile | Electron Microscopy Sciences | #10020 |
| UltraBed kit | Electron Microscopy Sciences | #14310 |
| 3,4-Dihydroxybenzonitrile, 98% | TCI | #D3352 |
| Silicone mold making kit | Let's Resin | #B096DKYT2C |
| **Algorithms** | | |
| EM thumbnail alignment | https://github.com/lichtman-lab/thumbnail_aligner | |
| EM fine alignment | https://github.com/lichtman-lab/mb_aligner | |
| X-ray tomography data processing | https://github.com/lichtman-lab/xradia_scripts | |

Methods

**Solutions.** The aCSF solution was composed of 125 mM NaCl, 26 mM $NaHCO_3$, 1.25 mM $NaH_2PO_4$, 2.5 mM KCl, 26 mM glucose, 1 mM $MgCl_2$ and 2 mM $CaCl_2$. $MgCl_2$ and $CaCl_2$ were added after the solution was bubbled with carbogen gas (95% $O_2$ and 5% $CO_2$) for 20 min. The aCSF solution was made fresh before each use and was used at room temperature.

The stock cacodylate buffer (2x) was composed of 300 mM sodium cacodylate, 8 mM $MgSO_4$, 4 mM $CaCl_2$, and was titrated with 1N HCl to a pH of 7.4. The divalent cations were added to preserve membrane integrity. (*43*)

**Tissue fixation.** We used neonatal (P0) and adult C57BL/6 mice for the protocol development and validation. All animal procedures were performed according to US National Institutes of Health guidelines and approved by the Committee on Animal Care at Harvard University.

P0 mice were anesthetized by hypothermia. The pups were placed in a latex glove and immersed up to the neck in crushed ice. After confirming unconsciousness, they were perfused transcardially for 5 minutes with the ice-cold fixative solution containing 2.5% glutaraldehyde, 2% paraformaldehyde, 2 mM $CaCl_2$, 4mM $MgSO_4$ and 150 mM sodium cacodylate buffer (pH 7.4) at a flow rate of 1 mL/min to match the cardiac output of the mouse at this age.

Adult mice were anesthetized with isoflurane inhalation until they no longer responded to paw pinch. The animals were transcardially perfused at a flow rate of 10 mL/min.

For conventional fixation that does not preserve ECS, room temperature fresh-made aCSF was perfused first for 2-3 minutes to clear blood. The liver should turn yellowish after blood removal. Then, the ice-cold fixative containing 2.5% glutaraldehyde, 2% paraformaldehyde, 2 mM $CaCl_2$, 4 mM $MgSO_4$, and 150 mM sodium cacodylate buffer (pH 7.4) was flowed for 5 minutes to induce fixation.

For ECS-preserved fixation, aCSF was perfused for 2-3 minutes to clear blood. The hypertonic solution containing 15% w/v mannitol in aCSF was perfused for 1 min to increase the permeability of blood-brain barrier (BBB). Then, the solution containing 5% w/v mannitol in aCSF was perfused for 5 minutes to restore the ECS. After this, the ice-cold fixative solution containing 4% w/v mannitol, 2.5% glutaraldehyde, 2% paraformaldehyde, 2 mM $CaCl_2$, 4 mM $MgSO_4$, and 150 mM sodium cacodylate buffer (pH 7.4) was flowed for 5 min to fix the whole brain with preserved ECS.

After perfusion, the brain was carefully removed from the skull to avoid mechanical damage, with fixative solution occasionally applied during the operation to keep the brain moist. The brain was placed into a biopsy Nylon mesh bag (VWR #10809-874) to protect it from mechanical irritation during staining and solution exchanges, and transferred to a 20 mL scintillation vial filled with fixative to post-fix at 4°C for 2 days under mild agitation before being processed for EM imaging. The fixed brains can be stored in the fixative for up to a month before use.

**ODeCO whole mouse brain staining.** We use adult mouse brain staining as an example to explain the detailed staining procedure. The step-by-step staining protocols for adult mouse brains, neonatal mouse brains and 1 mm brain slices are provided in the table below. We used a nutating rocker (VWR #82007-202) for gentle agitation, 20 mL glass scintillation vials for osmium staining, and 50 mL Falcon tubes for other steps.

1. Following fixation, the brain was transferred to a 50 mL tube containing cacodylate buffer (150 mM, with 2 mM $CaCl_2$ and 4 mM $MgSO_4$, pH 7.4) for thorough rinsing. The brain

was rinsed for 6 × 8 hours (2 days) at room temperature to ensure complete removal of the fixative. Any residual fixative may react with $OsO_4$ and affect tissue staining. (44)

2. Next, the brain was stained with 2% w/v $OsO_4$ buffered with 150 mM cacodylate (with 2 mM $CaCl_2$ and 4 mM $MgSO_4$) for 60 hours (2.5 days).
3. The osmicated brain was then rinsed with 150 mM cacodylate buffer (note: from this step, the cacodylate buffer no longer contains $Ca^{2+}$ and $Mg^{2+}$) for 8 × 12 hours (4 days).
4. After rinsing, the brain was destained with 2.5% w/v potassium ferricyanide in 150 mM cacodylate buffer. The destaining with ferricyanide was shielded from light to prevent side reactions. The solution was changed every 24 hours.
5. After 120 hours (5 days) of destaining, the brain was rinsed with 150 mM cacodylate buffer for 4 × 12 hours (2 days) and 150 mM NaCl for 4 × 12 hours (2 days).
6. Following rinsing, 2% w/v pyrogallol in 150 mM NaCl was applied. The tube was wrapped with aluminum foil to shield from light. The solution was changed after 24 hours.
7. After a total of 48 hours of conditioning with pyrogallol, the brain was rinsed with 150 mM NaCl for 8 × 12 hours (4 days). During the last rinse, the brain was moved to the refrigerator to cool down for the second osmication carried at 4°C.
8. After the rinse, the brain was transferred to a vial containing 2% w/v $OsO_4$ in 150 mM NaCl. The staining solution was changed when the staining was half-way done (3.5 days).
9. After a total of 7 days of second osmication, the brain was rinse with cold 150 mM NaCl at 4°C for 4 × 12 hours (2 days) and then moved to room temperature for rinsing with 150 mM NaCl for another 4 × 12 hours (2 days).
10. Subsequently, the brain was transferred to a gradual solution exchange container with an overflow hole connected to a vacuum trap (see Figure S10 for detailed setup). Over a 96-hours course, acetonitrile that was four times the volume of the dehydration container was injected, bringing the concentration to 98% acetonitrile. The brain was then transferred to a tube and rinsed with 100% acetonitrile for 4 × 6 hours (1 day) to remove any residual water.
11. For embedding, UltraBed low-viscosity epoxy resin was used. The resin was freshly prepared each time before use and degassed for about 10 minutes to remove dissolved gasses. Brain samples were infiltrated at room temperature with 25% resin:acetonitrile for 24 hours, 50% resin:acetonitrile for 24 hours, 75% resin:acetonitrile for 24 hours, 100% resin for 24 hours, and then fresh resin for another 24 hours.
12. The resin-infiltrated brain was removed from the Nylon mesh bag and placed into a customized silicone mold (see below for details on using 3D printed mouse brain models to create customized embedding molds). Resin-infiltrated samples was placed in a programmable oven (Yamato DKN-402C) held at 30°C for 12 hours, heated to 70°C over a 20-hour ramp, held at 70°C for 24 hours, and finally cooled down to 30°C over 20 hours.

**Table S2.** Summary of the staining steps of adult mouse brains, P0 mouse brains, and 1mm brain slices.

| Staining procedures | Solutions | Adult mouse brains | P0 mouse brains | 1mm brain slices |
|---|---|---|---|---|
| Rinse (r.t.) | 150 mM cacodylate buffer (with $Ca^{2+}$ and $Mg^{2+}$) | 6 × 8hr | 6 × 1.5hr | 4 × 0.5hr |
| 1st Osmication (r.t.) | 2% w/v $OsO_4$ in 150 mM cacodylate buffer (with $Ca^{2+}$ and $Mg^{2+}$) | 60hr | 8hr | 2hr |
| Rinse (r.t) | 150 mM cacodylate buffer | 8 × 12hr | 8 × 1.5hr | 6 × 0.5hr |
| Destaining (shield from light) | 2.5% w/v $K_3Fe(CN)_6$ in 150 mM cacodylate buffer | 120hr | 24hr | 6hr |
| Rinse (r.t.) | 150 mM cacodylate buffer transition to 150 mM NaCl solution | - 4 × 12hr with cacodylate buffer<br>- 4 × 12hr with NaCl solution | - 4 × 1.5hr with cacodylate buffer<br>- 4 × 1.5hr with NaCl solution | - 3 × 0.5hr with cacodylate buffer<br>- 3 × 0.5hr with NaCl solution |
| Conditioning (r.t., shield from light ) | 2% w/v pyrogallol in 150 mM NaCl | 48hr (change solution after 24hr) | 8hr | 2hr |
| Rinse (r.t. to 4°C) | 150 mM NaCl | 8 × 12hr (last rinse at 4°C) | 8 × 1.5hr | 6 × 0.5hr |
| 2nd Osmication (4°C) | 2% w/v $OsO_4$ in 150 mM NaCl | 168hr (4°C, change solution after 3.5 days) | 12hr (r.t.) | 3hr (r.t.) |
| Rinse (4°C to r.t.) | 150 mM NaCl | - 6 × 12hr (4°C)<br>- 2 × 12hr (r.t.) | 8 × 1.5hr (r.t.) | - 3 × 0.5hr with NaCl solution (r.t.)<br>- 3 × 0.5hr with $H_2O$ (r.t.) |
| Gradual dehydration for whole brains (r.t.) | Begin with 150 mM NaCl; Inject acetonitrile equivalent to 0.3 times the volume of the container during the first 24 | - 96hr gradual dehydration;<br>- 3 × 8hr with acetonitrile | - 20hr gradual dehydration<br>- 3 × 1hr with acetonitrile | - 0.5hr with 25% acetonitrile; |

| | | | | |
|---|---|---|---|---|
| | hours to reach a concentration of 25% acetonitrile/75% NaCl;<br>Replace the solution with a mixture of 25% acetonitrile/75% $H_2O$;<br>Inject acetonitrile equivalent to 3.7 times the volume of the container to reach a concentration of 98% acetonitrile;<br>Use 100% acetonitrile for the final rinse. | | | - 0.5hr with 50% acetonitrile;<br>- 0.5hr with 75% acetonitrile;<br>- 3 × 0.5hr with 100% acetonitrile |
| Resin infiltration (r.t.) | UltraBed diluted by acetonitrile | - 24hr with 25% resin;<br>- 24hr with 50% resin;<br>- 24hr with 75% resin;<br>- 24hr with 100% resin;<br>- 2 × 24hr with 100% resin | - 6hr with 25% resin;<br>- 12hr with 50% resin;<br>- 12hr with 75% resin;<br>- 12hr with 100% resin;<br>- 24hr with 100% resin | - 4hr with 25% resin;<br>- 8hr with 50% resin;<br>- 8hr with 75% resin;<br>- 8hr with 100% resin;<br>- 12hr with 100% resin |

**Sample screening.** Resin embedded brains were first screened by X-ray microCT (Zeiss Xradia 520 Versa) before cutting. X-ray microCT imaging was acquired by a 0.4× objective lens, with a tube voltage of 80kV and output power of 7W, exposure time at 10 seconds. Over a 360° rotation, 2401 projection images were acquired. Reconstruction was performed on Zeiss TXM3DViewer to verify the tissue quality. After screening, the crack-free block was then trimmed by a 3 mm UltraTrim diamond knife (Diatome) and ultramicrotome (UC6, Leica) to an elongated hexagon shape, with one diagonal oriented along the cutting direction. 30-40 nm thick serial sections at a selected depth were cut with a 6 mm wide Histo diamond knife (Diatome) that was equipped with the automated tape collection ultramicrotome (ATUM) at a cutting speed of 0.3 mm/s. Sections were collected onto the homemade carbon-coated Kapton tape. The tape was then cut into strips and mounted onto a 4-inch doped silicon wafer (University Wafers) via the double-coated carbon conductive tape (Ted Pella).

To enhance the signal, the wafer was plasma-treated for 30 seconds to increase its hydrophilicity of the Kapton tape and then post-stained with 4% uranyl acetate for 3 min, rinsed with deionized water for 30 seconds three times, air-dried, stained with 3% lead citrate solution (Leica Ultrostain II) for 3 min, rinsed with deionized water for 30 seconds three times, and air-dried. The post-stained wafer was mounted on a metal wafer holder with fiduciaries to target high-resolution imaging by the multi-beam scanning electron microscope (mSEM, Zeiss) and a low-resolution optical image 3.57 μm/pixel was acquired by Zeiss Axio Imager to record the position of each section relative to the fiduciaries. The wafer was then stored in a vacuum chamber before mSEM examination.

For P0 brains, whole sections selected at specific depth were imaged at 4nm/pixel. For the 3D dataset, a rectangular region of interest (ROI) was defined and superimposed onto each section in the optical image of that wafer using the Zen software package (Zeiss) to target high-resolution imaging in the mSEM. The selected sections and the pre-defined ROIs were imaged in the mSEM with a tile overlap of 10%, incident beam energy of 1.5 kV, dwell time 400 ns/pixel. Brightness and contrast for each ROI were set to maximize the dynamic range of the images acquired.

The acquired mSEM images were stitched by the SURF feature matching method *(45)*. An optimization step was followed to minimize the distances between the matched keypoints by finding an affine transformation for each image tile. After stitching, each 2D montage representing one section was aligned to its neighbors with a coarse-to-fine approach. The sections were first downsampled to micrometer resolution and coarsely aligned by an algorithm specifically designed for EM thumbnail alignment (https://github.com/lichtman-lab/thumbnail_aligner). After that, template matching was performed to obtain accurate corresponding points between neighboring sections with the help of the coarse transformations from the thumbnail alignment. Finally, a spring mesh system *(46)* was introduced to elastically morph each section according to the template matching results (https://github.com/lichtman-lab/mb_aligner).

**Time-lapse X-ray microCT imaging.** The time-lapse X-ray microscopy experiments were performed on a Zeiss Xradia 520 Versa X-ray microscope. The samples were wrapped in a biopsy Nylon mesh bag (VWR) and then immersed in a staining solution in a scintillation vial with the cap tightly sealed. The 3.7mL vials (EMS) were used for P0 brains and brain slices imaging, and the 10mL vials (EMS) for adult mouse brain imaging. The vial was then glued to the rotary stage in the X-ray recording chamber via double-sided tapes. Scans were acquired using the Scout-and-Scan Control System (Zeiss). Before each scan, adjust the sample position to the center of the field of view and set up the recipe according to the duration of each experiment. The sample was imaged using a 0.4× objective lens, at the tube voltage of 60kV and output power of 5W. The exposure time was set at 1.5 seconds for P0 mouse brains and brain slices, 2.5 seconds for adult mouse brains. For each tomographic scan, the sample was rotated 360° and 401 projection images were taken over one rotation. A tomographic scan took 25 min-35 min depending on the exposure time. If the experiment was longer than 12 hours, the experiment was suspended briefly every 12 hours to change the staining solution in the vial.

**X-ray data analysis.** The X-ray microCT reconstruction was output from Zeiss Reconstructor Scout-and-Scan software in Zeiss's proprietary *.txm* file format. The intensity values were normalized to cover the entire dynamic range of 16-bit images. To properly compare the X-ray absorptions between different images and to allow for other quantitative analyses, a Python API was developed to restore the image dynamic range to the original values and convert the images into the more standard *.nrrd* format (https://github.com/lichtman-lab/xradia_scripts). The 3d images were registered serially using Elastix toolbox (*47*) to generate the time series of cross-section views. After registration, we segmented each 3d image to measure the volume change using Huang's threshold method (*48*), followed by the Geodesic Active Contour method (*49*) to refine the boundaries. The segmentations were loaded to ImageJ and visually examined against the image volumes, and occasionally manual corrections were made if necessary. Finally, the 3D volumes were rendered with the Visualization Toolkit library (*50*).

**3D printing mouse brain models.** Custom-written Python code was used to extract the X-ray microCT volume from Zeiss .txm format to .nrrd format, as described above. The resulting nrrd volume was then loaded into ImageJ, where it was filtered with a Gaussian kernel to reduce noise and binarized using Otsu's method to create a binary mask. To ensure easy release during the downstream demolding process, each coronal section of the binary mask was dilated by half a millimeter and convexified. For tissue embedding, one handle was added to the anterior and posterior of the brain model as a gripping handle for cutting. Finally, the volumetric mask was loaded into ParaView, where it was meshed with the "Contour" filter and simplified with the "Decimate" filter. The resulting mesh was exported as a .stl file for 3d printing. VeroWhite was used as the 3D printing material on a Stratasys Object30 printer. The printed models were washed with detergent to remove unreacted residuals and placed at the bottom of a rectangular plastic container. The two-component silicone rubber kit was mixed in a 1:1 weight ratio and degassed with a vacuum oven to remove bubbles. The degassed silicone rubber was poured into the container to cover the brain models. The mold was left on the benchtop overnight to cure. The next day, the mold was removed from the container, and the mouse brain models were detached with the mold.
To create dummy mouse brains for use in the destaining test, serum-added agarose was warmed and poured into the mold (without handles). The solidified dummy brain was removed from the mold and stored in a fixative solution containing 2.5% glutaraldehyde, 2% paraformaldehyde, and 150 mM sodium cacodylate buffer (pH 7.4) before use.
For brain embedding, a thin layer of degassed fresh resin was added to the mold. Then, the resin-infiltrated mouse brains were carefully removed from the Nylon mesh bags and placed in the mold, ensuring no pressure was applied to squeeze the brain. A thin wooden stick can be used to adjust the brain to the correct position in the mold. Additional resin was added to fill the mold. The entire setup was then transferred to a programmable oven for curing.
**Method for automated segmentation and reconstruction.** The MapRecurse pipeline was used for the 3D segmentation of the P0 dataset. It includes fast-path steps (steps 1-4), paired merge-

error detection and correction steps (steps 5,7), and paired split-error detection and correction steps (steps 6, 8). The steps are as follows:

1. Prediction of membrane pixels using convolutional neural networks (CNNs) in the MATLAB-based environment "mEMbrain" (52).
2. Prediction of pixels straddling 2D skeletons.
3. Inferring 2D instance segmentation using local minima in the membrane probability map and region-growing watersheds.
4. Application of a set of fast object-linking algorithms: i) a matching algorithm based on 2D instances similar in shape and position across images; ii) linking overlapping objects based on the presence of overlapping pixels sufficiently far from detected membranes; iii) linking image-adjacent objects using predicted skeletal paths in the image; iv) matching objects between adjacent images if the predicted skeletons sufficiently match.
5. Merge errors were detected based on the rationale that neurons rarely include membrane-separated branches in contact with each other. The detected wrongly merged objects were corrected using an agglomerative approach that deconstructs complex erroneous objects with their agglomeration tree and reconstructs them iteratively while avoiding constructions that lead to large object self-contacts. Detecting such global features within the fast-path would not allow for globally optimizing the reconstruction and would require updating the agglomeration data structure, which was currently carried out once the linking step is completed.
6. Split errors were detected by identifying orphans that were not connected to large objects. Orphans were reconnected by rejoining the smallest number of edges from the priority queue of edge weights, eliminating the orphan property of the detected objects.

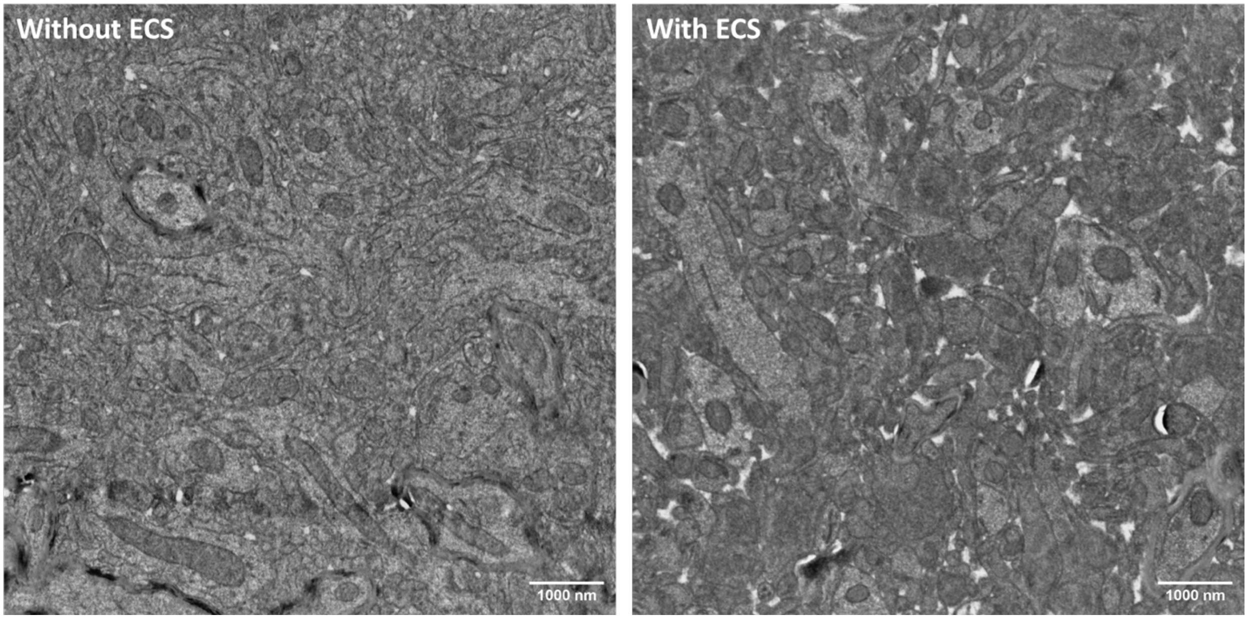


**Fig. S1.**
Overstaining of cytosolic proteins resulted in darkened cellular structures that are difficult to distinguish.

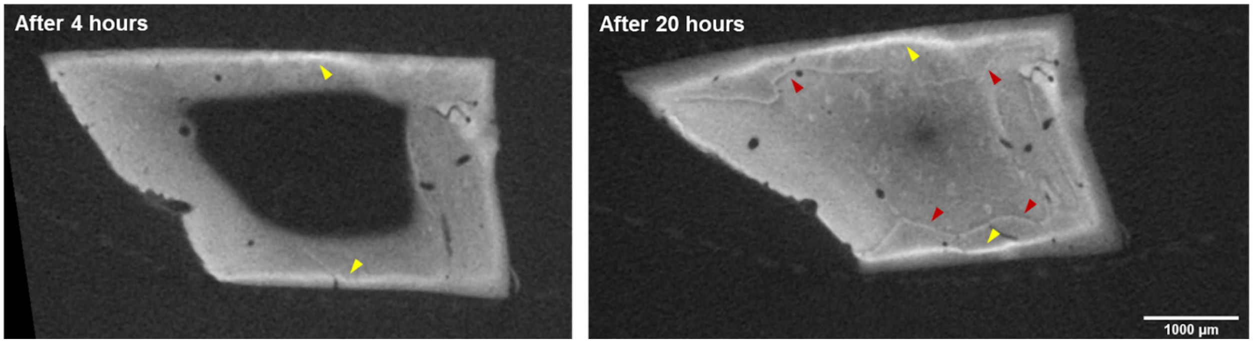


**Fig. S2.**
X-ray microCT of a 2.5mm-thick brain slab treated with reduced osmium (1 w/v% $OsO_4$ and 1.5 w/v% potassium ferrocyanide) in sodium cacodylate buffer. Cross-sectional view revealed osmium black precipitate approximately 150µm from the surface (a) after 4 hours of reaction, and (b) the occurrence of severe staining gradients and secondary precipitation after 20 hours of reaction. Yellow arrows indicate primary precipitation, while red arrows highlight secondary precipitation. Pixel values were not inverted to better display the precipitates which looked bright in the images.

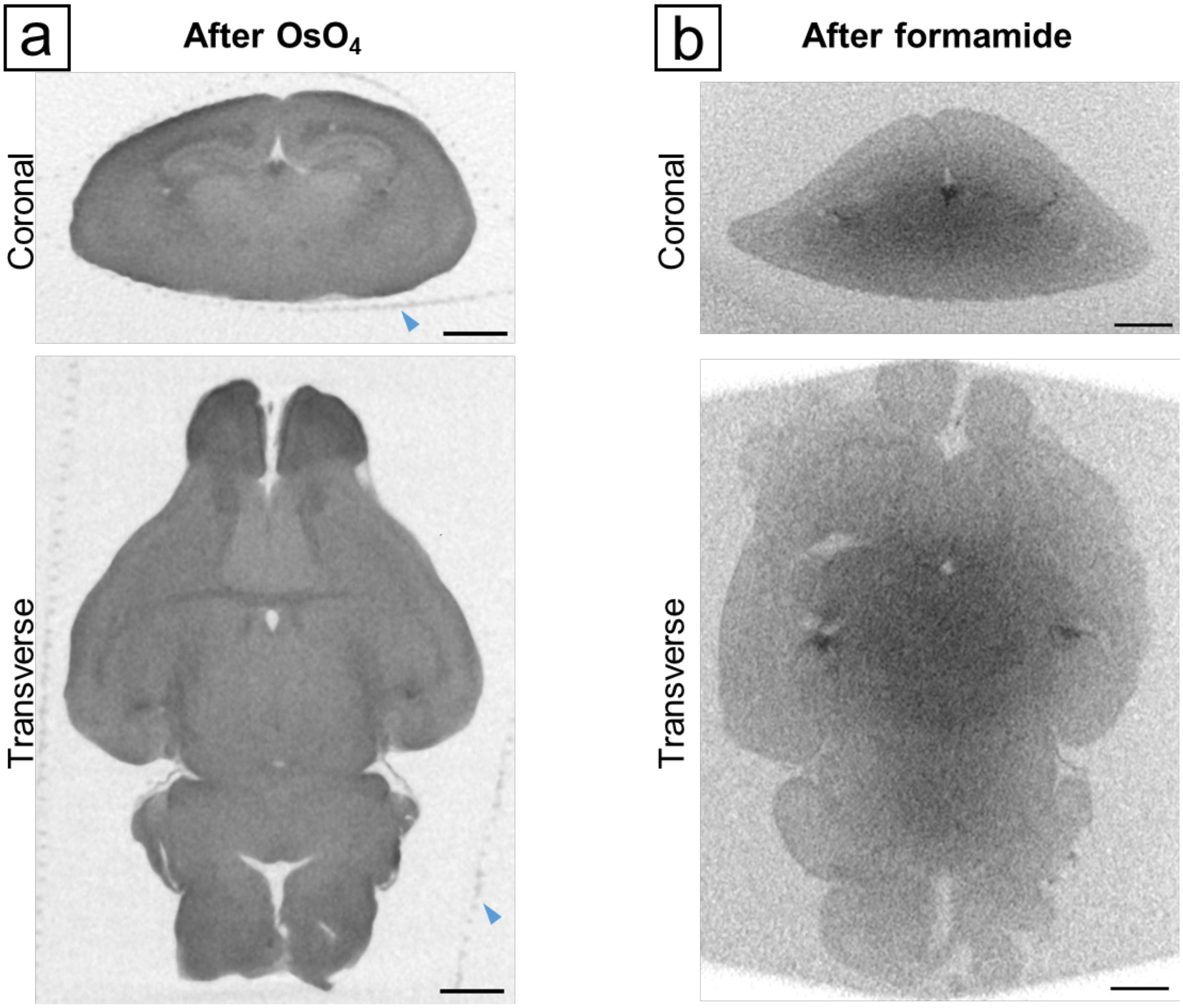


**Fig. S3.**
Formamide induced tissue softening. (a) A P0 mouse brain initially stained with 2 w/v% $OsO_4$ in sodium cacodylate buffer, followed by (b) treatment with 10 v/v% formamide and 2.5 w/v% potassium ferrocyanide in sodium cacodylate buffer. Formamide accelerated tissue destaining with ferrocyanide but also led to significant tissue distortion. Our brain specimens were enclosed in mesh nylon biopsy bags during staining to minimize mechanical disturbances from agitation and solution changes (blue arrow indicates nylon bag). Although typically benign, the formamide-induced tissue softening, resulting from protein crosslink dissociation, caused compression and flattening of the brain. Scale bars: 1 mm.

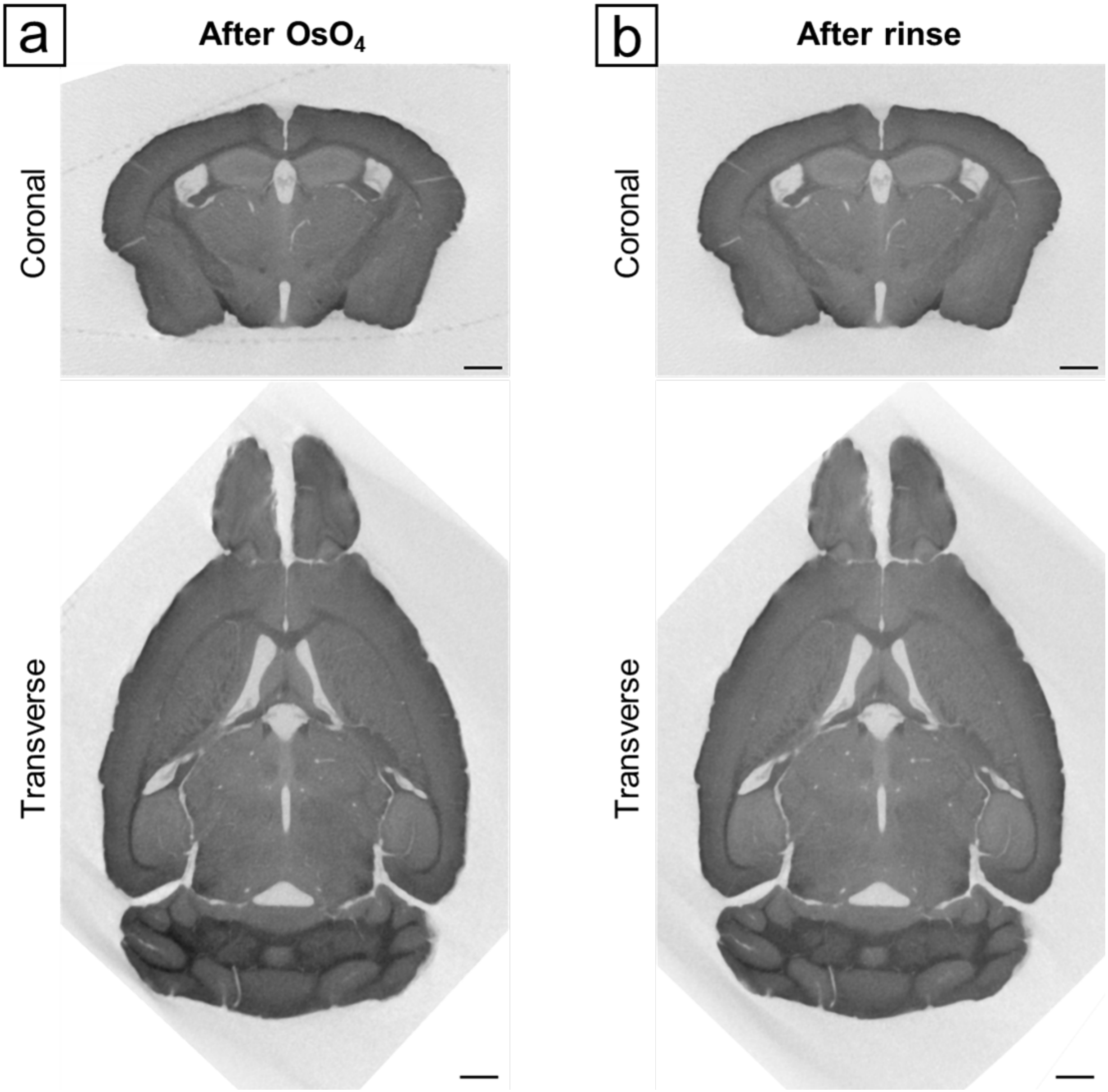


**Fig. S4.**

Thorough rinsing does not promote significant loss of osmium in tissue. (a) Adult mouse brain stained with 2 w/v% $OsO_4$ in sodium cacodylate buffer, followed by (b) 8 × 12-hour rinses with sodium cacodylate buffer. X-ray microCT in (b) was color normalized to (a). Comparisons revealed only a slight reduction in X-ray attenuation in (b), with a more pronounced decrease in gray matter but no noticeable change to the white matter (heavily myelinated regions). This is consistent with the expectation that rinsing primarily removes free-floating and loosely-bound osmium in cytosol without greatly affecting membrane-bound osmium. Scale bars: 1 mm.

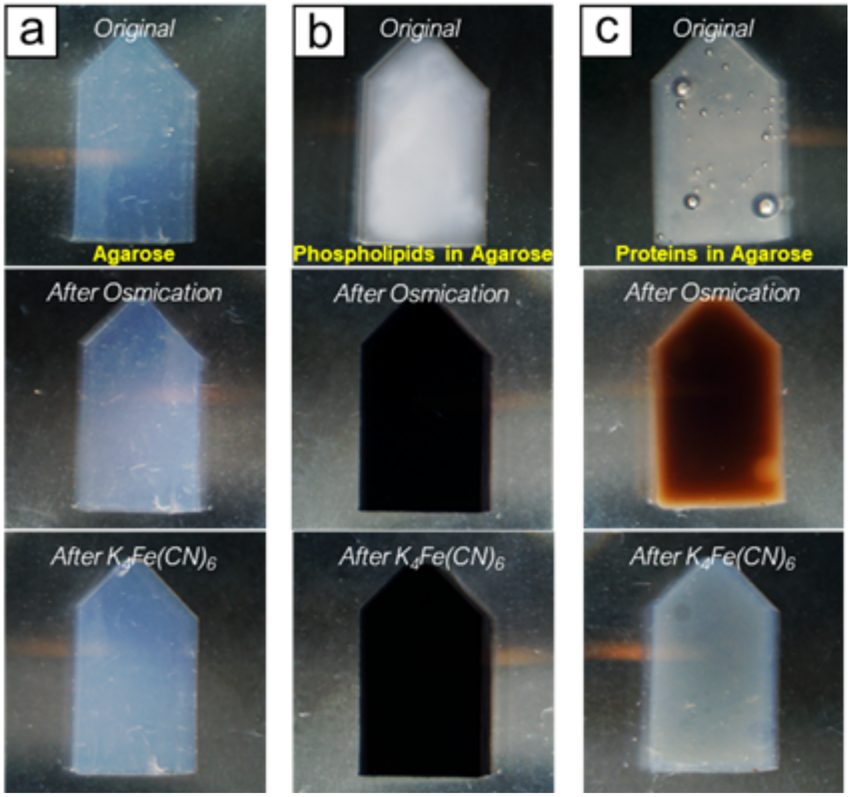


**Fig. S5.**
The agarose gel test shows that ferrocyanide mainly removes osmium bound to the proteins. The thin agarose gel (column a) was made from 5% w/v low-melting agarose (ThermoFisher); the phospholipids containing gel (column b) was made by adding 5% w/v Sphingomyelin (Santa Cruz Biotechnology) to the agarose gel; the protein gel (column c) was made by adding 10% v/v normal goat serum (ThermoFisher) to the agarose gel. (a) The pure agarose gel was translucent before the reactions and did not develop any color change after osmication and ferrocyanide rinse. (b) The phospholipids gel was opaque before the reactions. 1 w/v% buffered $OsO_4$ turned the gel to black immediately, and this color did not fade after rinsing with 2.5 w/v% buffered potassium ferrocyanide solution. (c) The protein gel was pale yellow before the reactions and turned to dark brown after staining in 1 w/v% buffered $OsO_4$. This color disappeared after rinsing with 2.5 w/v% buffered ferrocyanide solution for a day. However, the final block was slightly darker than the original one. For this test, the gel was about 2mm thick, thinner than the dummy brains, therefore could be effectively destained by ferrocyanide treatment.

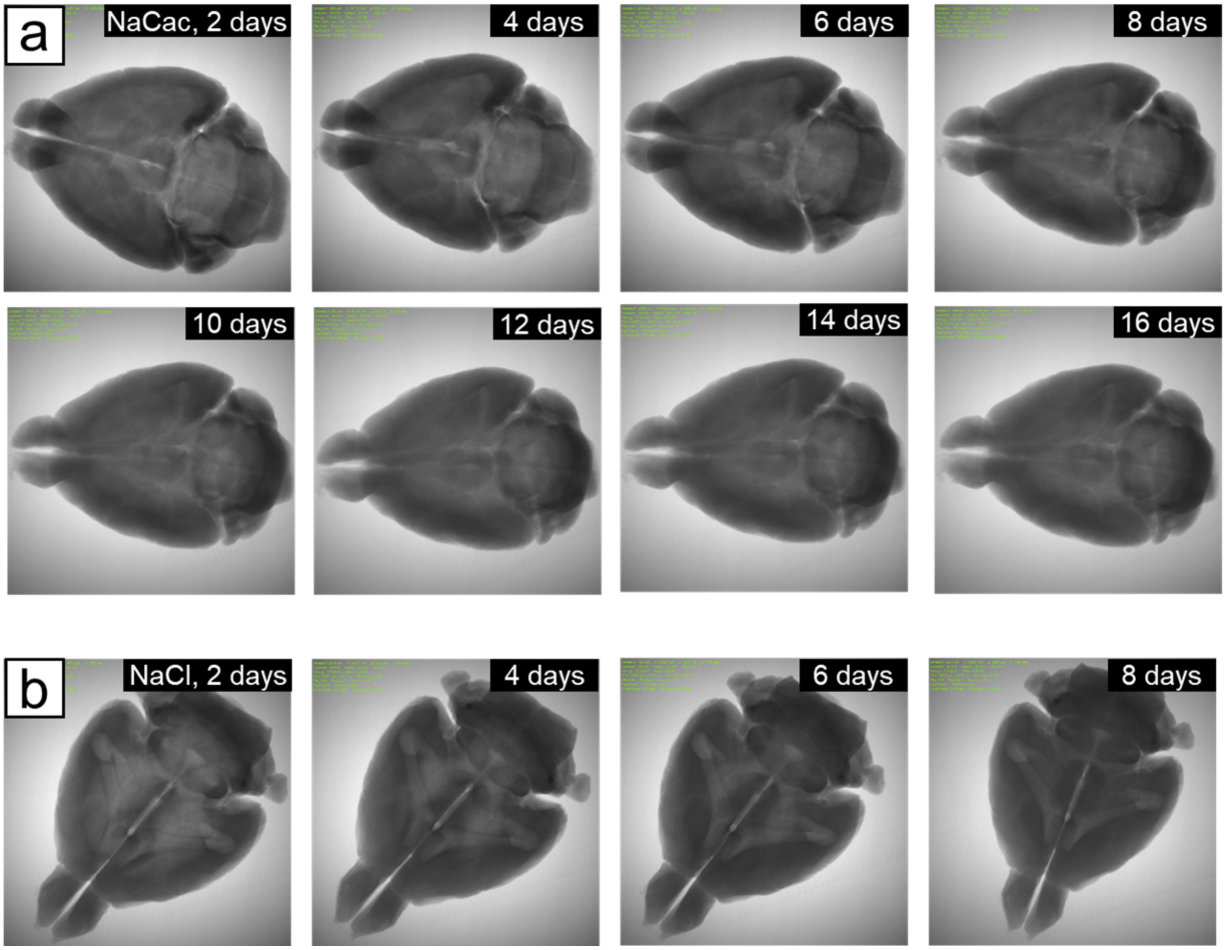


**Fig. S6.**
X-ray microCT projection comparison of two different reaction mediums for the second osmication performed at 4°C. (a) With sodium cacodylate as the reaction medium, the second osmication took over two weeks, and an under-stained core was still observed after 16 days. (b) Using sodium chloride, the mouse brain was fully stained after 8 days, with no under-stained core observed.

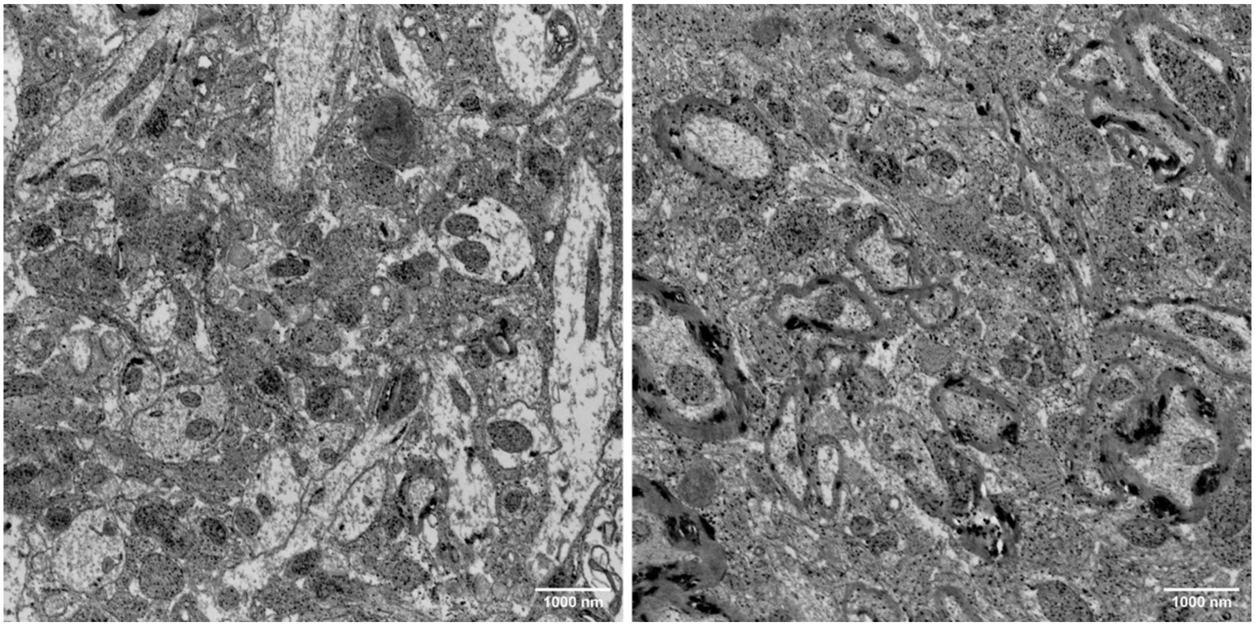


**Fig. S7.**
Sodium chloride is not an ideal reaction medium for the first osmication. When sodium chloride was used for the first osmication while keeping the rest of the protocol unchanged, the tissue was still stained; however, numerous dark precipitates appeared in the tissue.

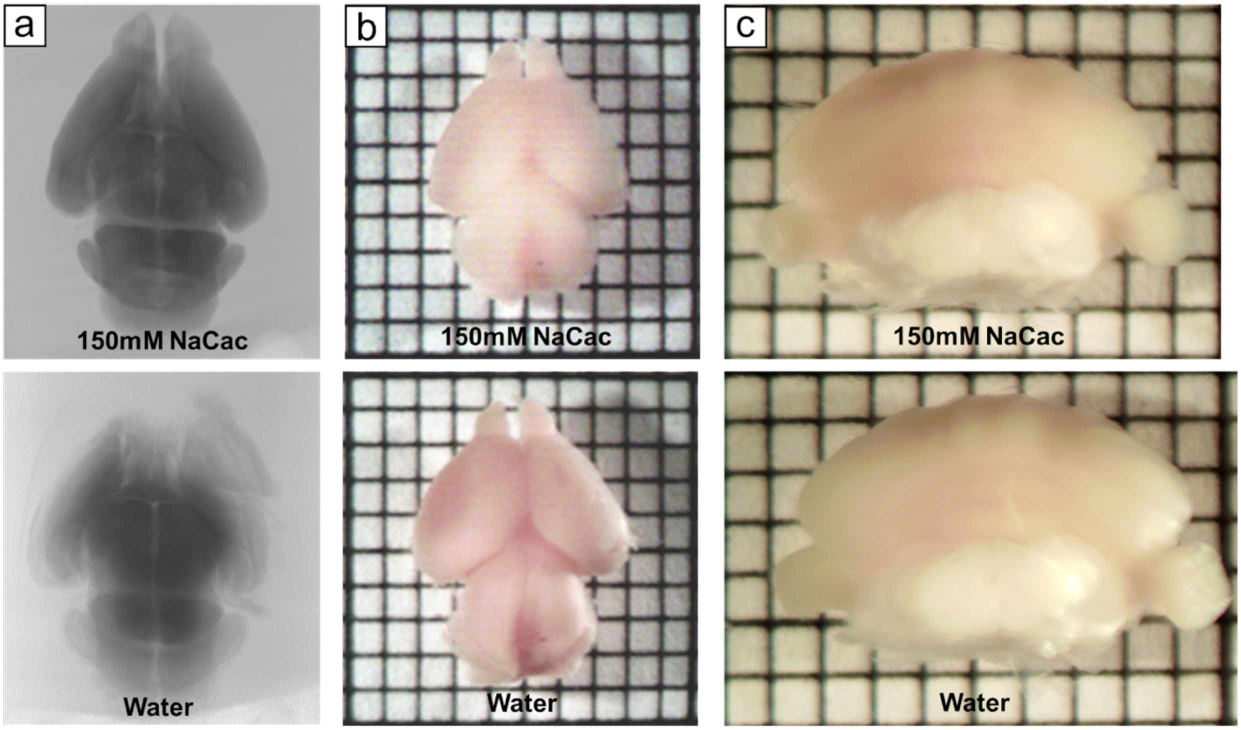


**Fig. S8.**
(a) X-ray microCT revealed that transferring an osmicated P0 mouse brain from buffer to water resulted in immediate tissue expansion, causing the entire brain to disintegrate. The transition of a chemically fixed (b) P0 mouse brain and (c) adult mouse cerebellum to water also led to perceivable expansion.

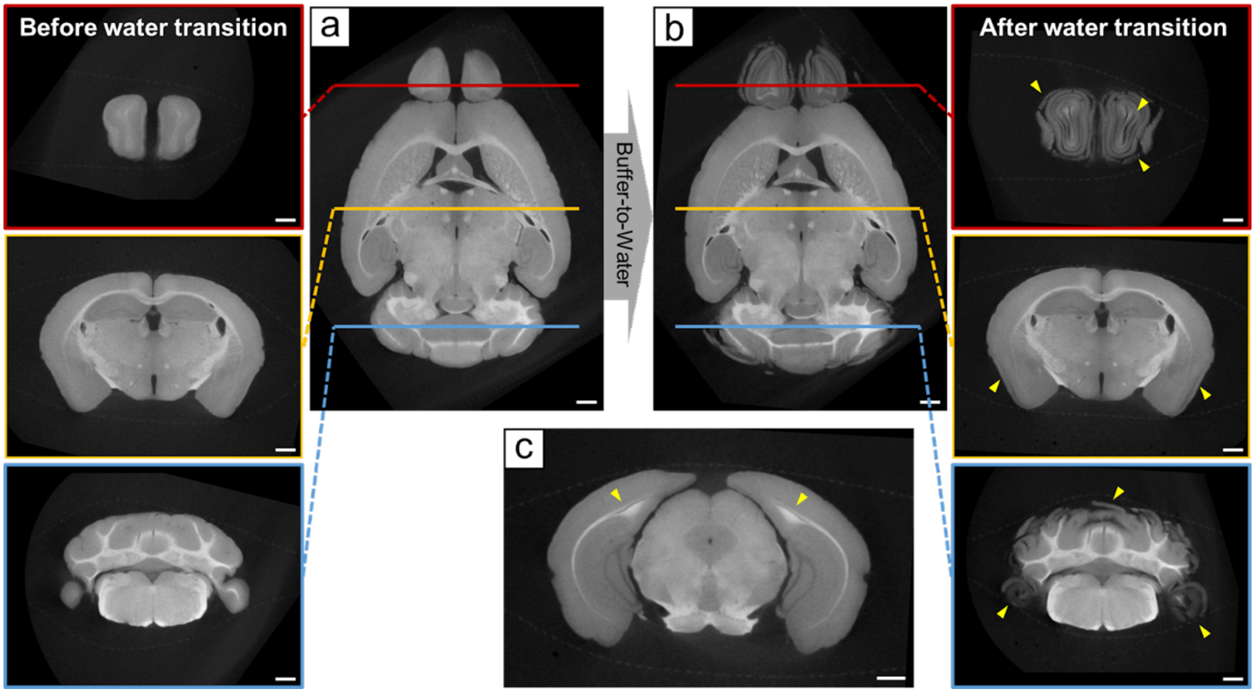


**Fig. S9.**
Replication of Song's whole-brain staining protocol (*12*, *51*) was performed on three adult mouse brains (two with conventional fixation and one with ECS-preserved fixation; this figure showcased one sample with conventional fixation). All the brain specimens noticeably expanded and became brittle after the transition from buffer to water. X-ray microCT scans of the specimens (pixels were not inverted), taken after each step, revealed that the transition from buffer to water led to severe tissue fragmentation mainly in the anterior and posterior regions of the brains. (a) After the second osmium tetroxide incubation, intended to increase tissue stability and tolerance to osmotic stress, the brain was rinsed with cacodylate buffer. At this stage, the brain remained intact and free of cracks. (b) Subsequent to the buffer rinse, the brain was rinsed with water, which caused the crumbling of the olfactory bulb and the cerebellum (indicated by yellow arrows). (c) The central part of the brain exhibited minimal damage, with only the subcortical, highly myelinated regions showing a few cracks (indicated by yellow arrows), as described in Song's paper. We believe both tissue fragmentation and cracking were caused by the immediate volume expansion associated with the buffer-to-water transition and need to be addressed accordingly.

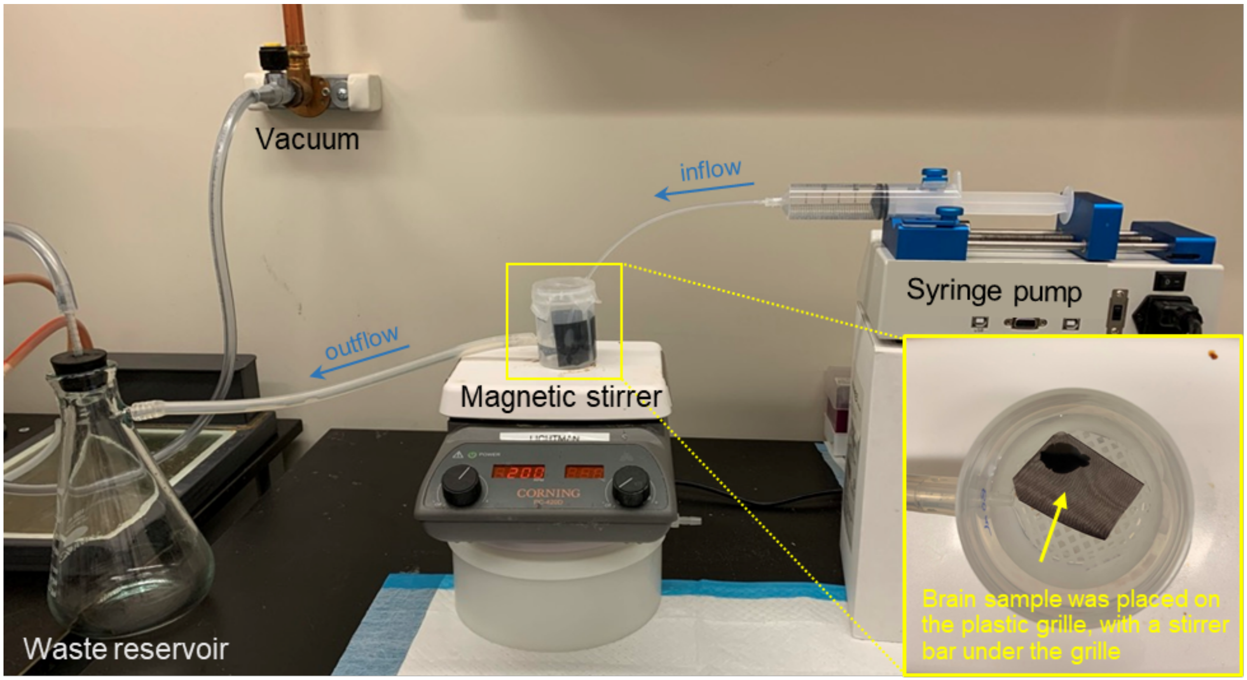


**Fig. S10.**
Gradual solution exchange device setup. The brain specimen wrapped in a biopsy Nylon mesh bag was placed on a plastic grill above the stirrer bar. For one brain, a container holding more than 40 mL solution was used. The container was filled with 150 mM NaCl solution up to the overflow hole on the sidewall (i.e., 40 mL). Acetonitrile was slowly injected into the container using a syringe pump (Fusion Touch) at the preset flow rate and continuously mixed with the solution by the spinning stirrer bar. The overflowing solution was slowly drained into a glass vacuum trap with a low vacuum applied. The container was sealed with parafilm to prevent excessive evaporation. The first 12 mL of acetonitrile was injected at a rate of 0.5 mL/hr. After 24 hours, the solution in the container reached 25% acetonitrile/75% NaCl. The injection was paused, and the solution was replaced with a 25% acetonitrile/75% DI-$H_2O$ mixture to prevent the precipitation of NaCl in a non-polar solvent. The injection then resumed at a rate of 2 mL/hr, with 148 mL of acetonitrile being injected over 74 hours. A total of 160 mL of acetonitrile, four times the container's volume, brought the final concentration to 98% acetonitrile. The brain was then transferred to pure acetonitrile for a final rinse. (Inset) The solution was continuously mixed by a magnetic stirrer bar at the bottom of the vessel.

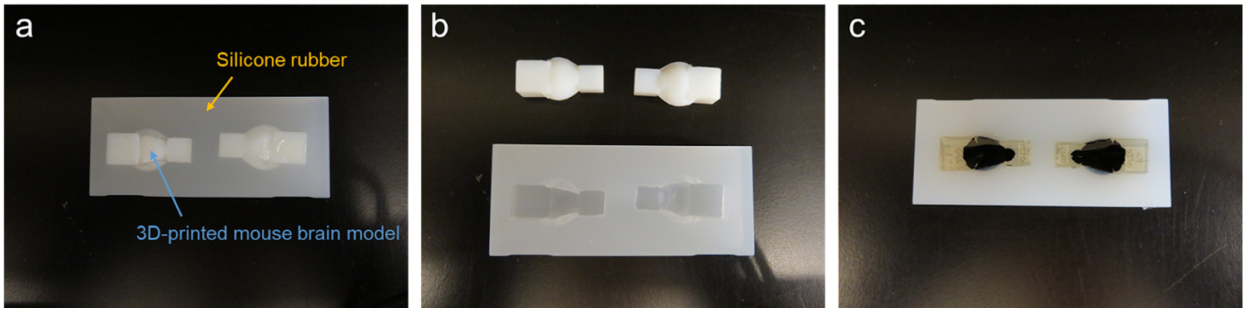


**Fig. S11.**
Customized silicone mold for whole-brain embedding to minimize surrounding resin. (a) 3D-printed mouse brain models, with two gripping handles in the anterior and posterior, were used with silicone rubber to create the embedding mold. (b) After silicone solidification overnight, the plastic mouse brain models were carefully detached from the mold, ensuring the silicone did not peel off with them. (c) Resin-infiltrated mouse brains fit nicely into the mold with gentle nudges.

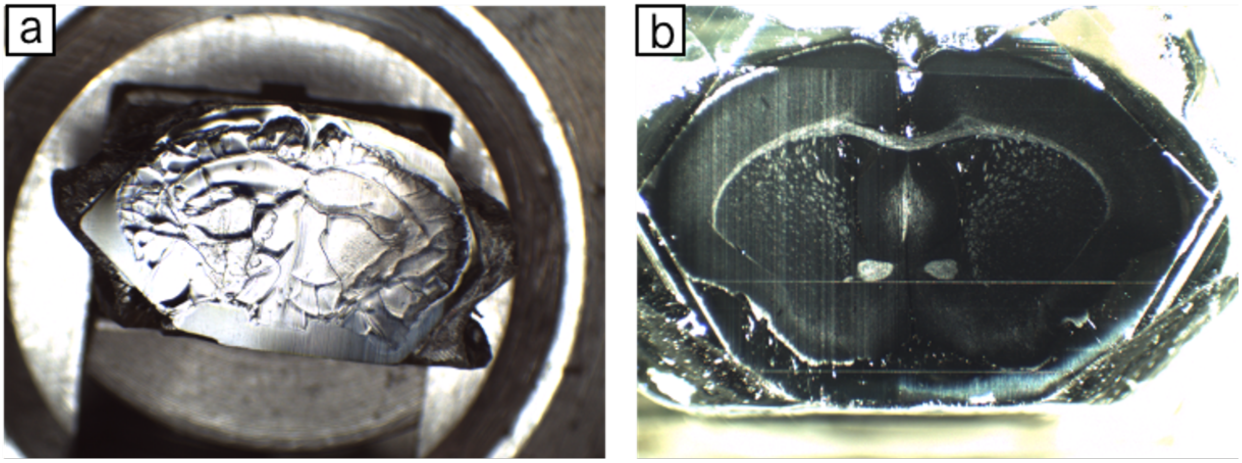


**Fig. S12.**
(a) Internal residual stress built up in the tissue during resin curing caused cracks to form when the block was cut open. (b) Customized molds and ramped heating and cooling minimized internal residual stress, and no cracks were observed when the block was cut open for tissue screening.

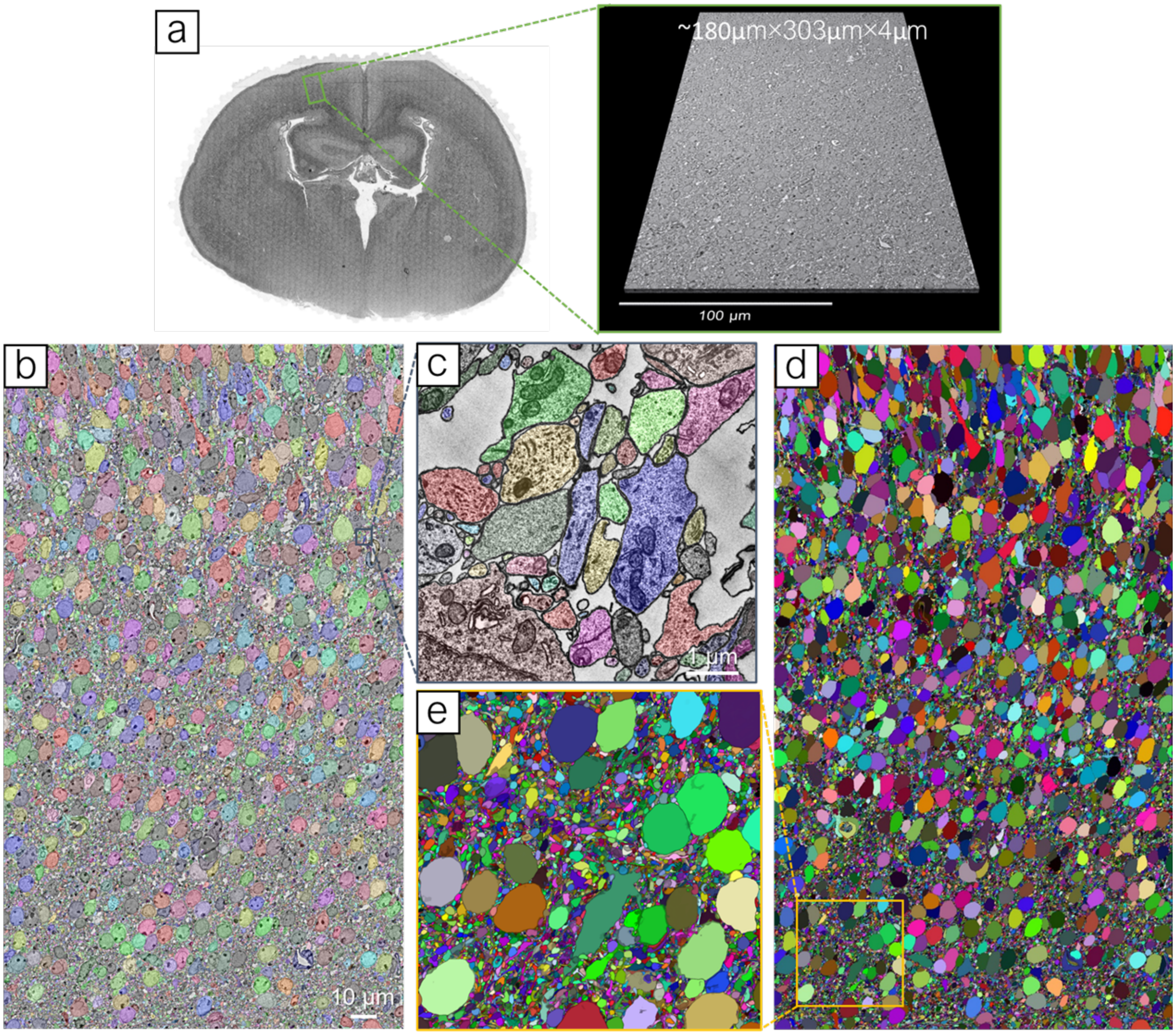


**Fig. S13.**
Segmentability assessment of an ODeCO-stained mouse brain. (a) 109 consecutive coronal sections of approximately 35nm each were collected from a stained P0 mouse brain. Using multiSEM, we acquired an image stack of ~180μm×303μm×4μm with x-y resolution of 4nm×4nm from those consecutive coronal sections. This stack covered layers II/III through VI of the secondary motor cortex in the neonatal mouse brain. (b) A colorized section from the image stack demonstrated the results of the auto-segmentation. (c) A magnified view of a small region highlighted the high quality of the auto-segmentation. (d) A dense 3D reconstruction of the entire stack and (e) a zoomed-in view of a small region displays that both somata and finest neuropils were accurately reconstructed.
